# The *Mycobacterium tuberculosis* Pup-proteasome system regulates nitrate metabolism through an essential protein quality control system

**DOI:** 10.1101/470989

**Authors:** Samuel H. Becker, Jordan B. Jastrab, Avantika Dhabaria, Catherine T. Chaton, Jeffrey S. Rush, Konstantin V. Korotkov, Beatrix Ueberheide, K. Heran Darwin

## Abstract

The human pathogen *Mycobacterium tuberculosis* (*M. tuberculosis*) encodes a proteasome that carries out regulated degradation of bacterial proteins. It has been proposed that the proteasome contributes to nitrogen metabolism in *M. tuberculosis*, although this hypothesis had not been tested. Upon assessing *M. tuberculosis* growth in several nitrogen sources, we found that a mutant strain lacking the *Mycobacterium* proteasomal activator Mpa was unable to use nitrate as a sole nitrogen source due to a specific failure in the pathway of nitrate reduction to ammonium. We found that the robust activity by the nitrite reductase complex NirBD depended on expression of the *groEL*/*groES* chaperonin genes, which are regulated by the repressor HrcA. We identified HrcA as a likely proteasome substrate, and propose that the degradation of HrcA is required for the full expression of chaperonin genes. Furthermore, our data suggest that degradation of HrcA, along with numerous other proteasome substrates, is enhanced during growth in nitrate to facilitate the de-repression of the chaperonin genes. Importantly, growth in nitrate is the first example of a specific condition that reduces the steady-state levels of numerous proteasome substrates in *M. tuberculosis*.

**SIGNIFICANCE STATEMENT:** The proteasome is required for the full virulence of *M. tuberculosis*. However, the extent of its role as a regulator of bacterial physiology remains unclear. In this work, we demonstrate a novel function of the proteasome system in maintaining the expression of essential chaperonin genes. This activity by the proteasome is required for *M. tuberculosis* to use nitrate as a nitrogen source. Furthermore, we identified a specific growth condition that robustly decreases the abundance of pupylated proteins. This observation strongly suggests the presence of a yet-to-be-determined mechanism of control over the Pup-proteasome system in *M. tuberculosis* that is induced in nitrate.

## INTRODUCTION

*Mycobacterium tuberculosis* (*M. tuberculosis*), the causative agent of the human disease tuberculosis, encodes a proteasome that is essential for its lethality in mice (1, 2). The central component of all proteasomes is a 28-subunit complex of four stacked rings known as the 20S core particle (20S CP). In *M. tuberculosis*, two identical outer rings, each composed of seven α-subunits (PrcA) serve as a gated entryway for protein substrates, and two identical inner rings, composed of a total of 14 β-subunits (PrcB), form the catalytic active sites of the protease (1, 3-5). While essential in eukaryotes and archaea, proteasomes are found only in a subset of bacteria primarily belonging to the *Actinomycetales* and *Nitrospirales* orders, and are not always essential for bacterial viability (6, 7).

In eukaryotes and bacteria, proteasomes carry out the regulated proteolysis of specific cellular substrates. Interest in the *M. tuberculosis* proteasome emerged after a screen for mutations that rendered this bacterial species sensitive to nitric oxide (NO), a host-derived molecule that is critical for controlling *M. tuberculosis* growth in mice (8), identified mutations in genes linked to *prcBA*. Over the years, it was determined that some proteasome substrates in *M. tuberculosis* are covalently modified with a small protein called Pup (<u>p</u>rokaryotic <u>u</u>biquitin-like <u>p</u>rotein) by a dedicated ligase, PafA (<u>p</u>roteasome <u>a</u>ccessory <u>f</u>actor <u>A</u>) (9-11). These pupylated proteins are recognized by a proteasomal activator, Mpa (<u>m</u>ycobacterial <u>p</u>roteasome <u>A</u>TPase, also known as ARC), which uses ATP hydrolysis to power the unfolding and delivery of proteins into 20S CPs for degradation (1, 12). Pup can also be removed from substrates by an enzyme called Dop (<u>d</u>eamidase <u>o</u>f <u>P</u>up) (13, 14), as well as by PafA (15). Collectively, Dop, PafA, Pup, Mpa and 20S CPs constitute the core “Pup-proteasome system” (PPS). At least sixty *M. tuberculosis* proteins are currently known to be pupylation substrates (9, 16, 17), while studies performed in other Pup-bearing bacteria, including *Mycobacterium smegmatis* (*M. smegmatis*), have identified hundreds of additional potential targets of pupylation (18-21). Of note, many pupylated proteins in *M. tuberculosis* are not degraded under routine culture conditions for reasons that remain unknown (16). This observation suggests pupylation may not immediately send proteins to the proteasome and could possibly serve a non-degradative regulatory role, as is observed in *Corynebacteria* (22).

In addition to being highly sensitive to NO *in vitro*, PPS mutants are highly attenuated for virulence in mouse infection models (2, 12, 23). The failure to degrade a single pupylated substrate, Log, is responsible for the NO hypersensitivity phenotype of a PPS (*mpa*) mutant. However, while genetic disruption of *log* completely restores NO resistance to an *mpa* strain *in vitro*, it does not fully rescue the virulence defect of this strain in mice (17). Therefore, there are likely to be other components of *M. tuberculosis* physiology whose regulation by the PPS is important for establishing lethal infections.

In addition to its central role in the post-translational regulation of various cellular pathways, an essential function of the eukaryotic proteasome is to maintain nutrient homeostasis by recycling amino acids (24, 25). In light of this observation, there has been interest in the question of whether or not the proteasome has a similar function in bacteria. Studies in *M. smegmatis* suggest that pupylation is required to maintain nitrogen homeostasis. Deletion of *pup* renders *M. smegmatis* more sensitive to nitrogen starvation (26), during which several enzymes involved in nitrogen metabolism are pupylated (21). In *M. tuberculosis*, amino acids serve as the primary nitrogen donors for most anabolic processes (27, 28). Additionally, optimal *M. tuberculosis* growth, both *in vitro* and *in vivo*, requires the uptake of exogenous amino acids as a nitrogen source (29-32). It has therefore been hypothesized that the products of bulk proteolysis by the *M. tuberculosis* proteasome could be an important source of nitrogen under nutrient-limiting conditions. For this reason, we sought to determine if the *M. tuberculosis* proteasome contributed to nitrogen metabolism. Contrary to what was observed in *M. smegmatis*, we found that proteasomal degradation did not provide a survival advantage to *M. tuberculosis* during nitrogen starvation. However, we discovered that the proteasome was essential for the ability of *M. tuberculosis* to use nitrate as a nitrogen source. Through a genetic suppressor screen, we identified a putative PPS substrate whose inactivation rescued the ability of an *M. tuberculosis* PPS mutant to assimilate nitrogen from nitrate. Our data revealed an essential role for the PPS to facilitate the activity of nitrite reductase, possibly in two different ways, during growth in nitrate. Finally, we found that the abundance of the pupylome decreased when *M. tuberculosis* was grown in nitrate, the first condition known to alter proteasome substrate levels in *M. tuberculosis*.

## RESULTS

### The *M. tuberculosis* proteasome does not provide a survival advantage during nitrogen starvation *in vitro*

It has been previously reported that an *M. smegmatis pup* (also known as *prcS* in *M. smegmatis*) or *prcBA* mutant cannot survive as well as a wild type (WT) strain during several weeks of nitrogen starvation; however, the phenotypes of these mutants are almost fully complemented by *pup* alone, suggesting that proteasomal degradation itself may have a minor role in *M. smegmatis* nitrogen metabolism. Nonetheless, it was proposed that the proteasome supported bacterial survival during nitrogen starvation by recycling amino acids (26). We therefore sought to test whether or not proteasomal degradation contributed to *M. tuberculosis* survival during nitrogen starvation. We incubated WT, ∆*mpa::hyg* ("*mpa*") and ∆*prcBA::hyg* ("*prcBA*") strains (see *SI Appendix*, Table S1) in Proskauer-Beck (PB) minimal medium lacking any nitrogen source and measured bacterial survival over time. In contrast to what is observed in *M. smegmatis*, we found that the WT strain had no survival advantage over the PPS mutant strains during three weeks of nitrogen starvation (Fig. 1A). Thus, amino acid recycling by the proteasome did not significantly contribute to nitrogen homeostasis in *M. tuberculosis* in our experiments. However, we cannot rule out a role for the proteasome in recycling amino acids under other conditions.

**Figure 1.**
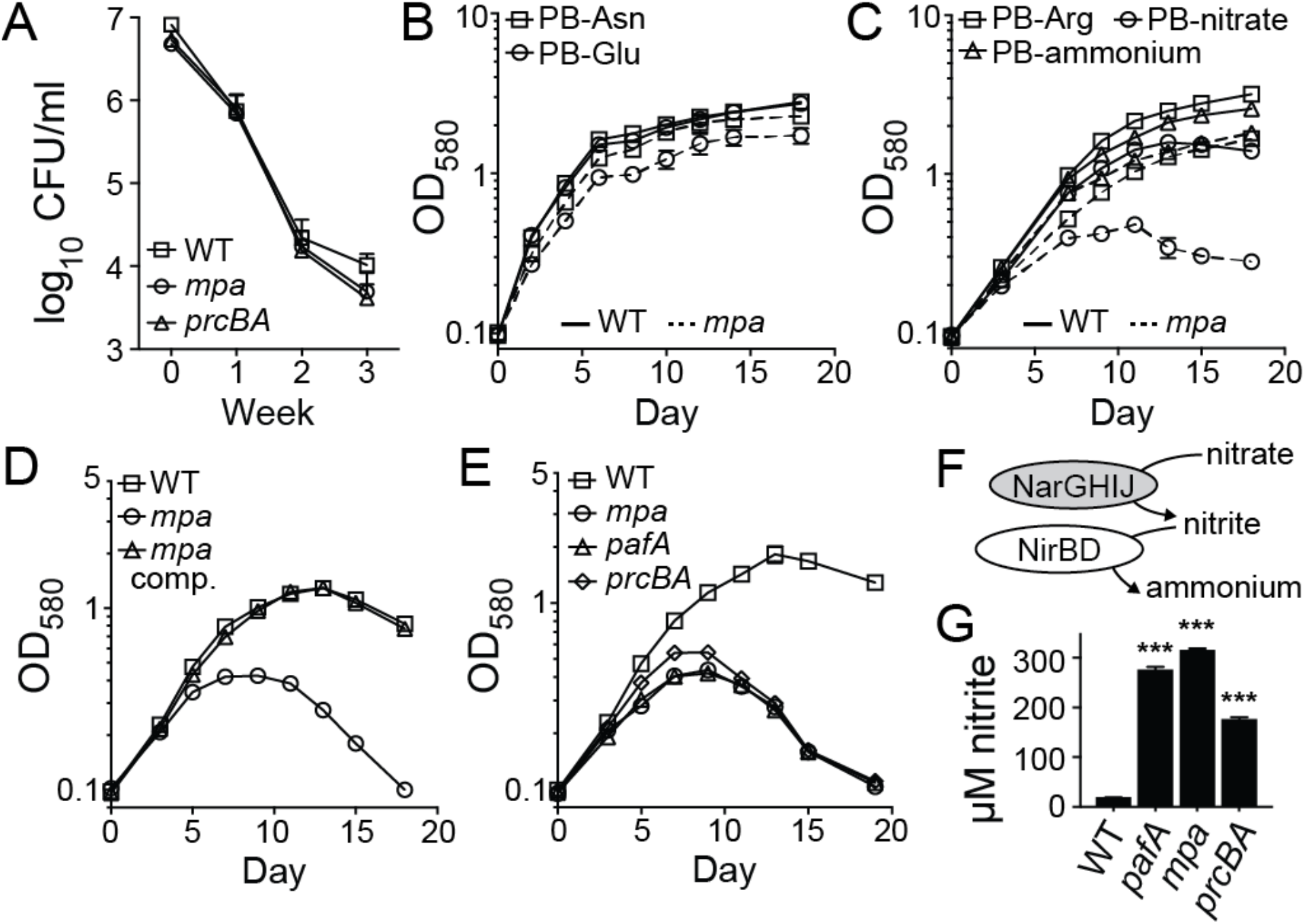
The *M. tuberculosis* Pup-proteasome system (PPS) is required for growth in nitrate. (A) The PPS does not promote survival of *M. tuberculosis* during complete nitrogen starvation. Survival of *M. tuberculosis* wild type (WT), *mpa* (MHD149), and *prcBA* strains was measured by number of colony-forming units (CFU) per ml of culture at the indicated time points. At week three, the fold change in CFU from input was determined to be statistically insignificant (*p* > 0.05) for *mpa* and *prcBA* strains compared to the WT strain. Experiment represents data from six replicate cutures. (B) The PPS is not essential for growth of *M. tuberculosis* in ideal nitrogen sources. Growth *of M. tuberculosis* strains in Proskauer-Beck (PB) minimal media supplemented with single nitrogen sources asparagine (PB-Asn) or glutamate (PB-Glu) was measured by optical density at 580 nm (OD_580_). (C) An intact PPS is essential for *M. tuberculosis* growth when provided nitrate as the sole nitrogen source. *M. tuberculosis* strains were grown in PB supplemented with arginine (PB-Arg), nitrate (PB-nitrate), or ammonium (PB-ammonium). (D) Complementation of the *mpa* mutant growth defect in PB-nitrate. (E) Pupylation and proteasomal degradation are required for *M. tuberculosis* nitrate utilization, as assessed by the growth of *pafA* (MHD2), *mpa* (MHD5), and *prcBA* strains in PB-nitrate. (F) Schematic of the *M. tuberculosis* enzymes that catalyze reduction of nitrate to ammonium (33, 34). (G) PPS mutants (as in E) secrete excess nitrite into culture supernatants during growth in PB-nitrate. Experiments in (B) through (E) and (G) each contain data from three replicate cultures. ***, *p* < 0.001.

### *M. tuberculosis* requires the PPS in order to use nitrate as a nitrogen source

Following our observation that *M. tuberculosis* proteasome-defective strains did not have a survival disadvantage during nitrogen starvation, we next determined if the PPS was required for growth in a specific nitrogen source. *M. tuberculosis* can use both organic and inorganic sources of nitrogen, although asparagine and glutamate support growth most effectively *in vitro* (29). We compared the growth of WT and *mpa* strains in PB media supplemented with asparagine or glutamate, and found that the *mpa* mutant had a minor growth defect compared to the WT strain (Fig. 1B). When the same strains were provided the sub-optimal nitrogen sources arginine, ammonium, or nitrate, bacterial growth was predictably slower for both strains; remarkably, however, growth of the *mpa* mutant was almost completely abrogated in nitrate compared to the WT strain (Fig. 1C). A single copy of *mpa* integrated on the chromosome restored growth of the *mpa* strain in nitrate (Fig. 1D).

To determine if the inability of an *mpa* mutant to use nitrate was specifically related to a failure to degrade pupylated proteins, we assessed the growth of a *pafA* mutant (*pafA*::MycoMarT7) and the *prcBA* strain. Both mutants were attenuated for growth similarly to an *mpa* mutant in PB-nitrate, demonstrating that both pupylation by PafA and proteolysis by 20S CPs were required for using nitrate as a sole nitrogen source (Fig. 1E).

*M. tuberculosis* uses a highly conserved pathway for nitrogen assimilation from nitrate (Fig. 1F). Once imported into the cell, nitrate is reduced to nitrite by the NarGHIJ nitrate reductase complex (33). Nitrite is then reduced to ammonium by the nitrite reductase complex NirBD (34). Finally, ammonium is incorporated into glutamate and glutamine, which comprise the major intracellular nitrogen pool (27). Notably, *M. tuberculosis* secretes into its extracellular space any nitrite that cannot be immediately reduced to ammonium (34).

We hypothesized that the inability of PPS mutants to productively grow in PB-nitrate was caused by a failure of one or more reactions within nitrate catabolism. Upon growing *M. tuberculosis* in PB-nitrate, we discovered that supernatants of *pafA*, *mpa*, and *prcBA* mutant cultures contained ten- to fifteen-fold higher concentrations of nitrite than those of a WT strain (Fig. 1G). This result suggested that these mutants, while capable of importing and reducing nitrate, were unable to reduce nitrite to ammonium, causing the secretion of excess nitrite. Further supporting this model, an *mpa* mutant strain was capable of growing in PB-ammonium, which bypasses the requirement of nitrite reduction, nearly as well as the WT strain (Fig. 1C).

In *M. tuberculosis*, the nitrite reductase complex is encoded by the *nirBD* (Rv0252-Rv0253) operon. An *M. tuberculosis nirBD* mutant is unable to grow when nitrate is provided as the single nitrogen source, implicating NirBD as the only nitrite reductase in *M. tuberculosis* (34). Therefore, we hypothesized that degradation of one or more pupylated proteins is required for the *in vivo* activity of NirBD.

### Suppressor mutations in *hrcA* or *nadD* restore growth of an *mpa* mutant in nitrate

Most of an *mpa* mutant culture ultimately dies upon extended incubation in PB-nitrate (Fig. 1A), an observation that provided a powerful phenotype to screen for suppressor mutations that might identify specific substrates of the PPS whose degradation is necessary for NirBD activity. We previously generated a transposon mutant library of an *M. tuberculosis mpa* strain, consisting of approximately 72,000 unique double-mutant clones (17). To enrich for mutants with a suppressor phenotype, we incubated this library in PB-nitrate for four to five weeks, until cultures were turbid. Surviving bacteria were further expanded in rich media and subjected to a second round of incubation in PB-nitrate. We isolated 16 clones from six independent pools and tested them individually for growth in PB-nitrate. Interestingly, while all of the suppressor mutants grew more productively than the parental *mpa* strain, no mutant grew as well as the WT strain (*SI Appendix*, Fig. S1A).

To identify the suppressor mutations, we cloned DNA containing the transposon insertion from each of the 16 isolates. We identified two strains with unique transposon insertions in the coding region of *hrcA* (Rv2374c). The remaining suppressor strains contained transposon insertions in different operons, with no obvious functional connections. We therefore suspected that these strains had additional, spontaneous mutations, possibly in *hrcA*. We PCR-amplified and sequenced the *hrcA* gene in the remaining mutants and discovered that five more strains had point mutations in *hrcA*. We next used whole-genome sequencing to identify mutations in the remaining nine suppressor mutants; seven strains had mutations in either the promoter or coding region of *nadD* (Rv2421c). Importantly, all of these suppressor mutations were the result of independent events (see *SI Appendix*, Table S1).

### Expression of chaperonin genes promotes growth in nitrate

Seven of the suppressor strains had mutations in *hrcA*, which encodes a transcriptional repressor that is conserved in many bacterial species as a regulator of molecular chaperones of the Hsp60 family [reviewed in (35)]. In *M. tuberculosis*, HrcA directly represses four genes, including three chaperonin-encoding genes *groES* (Rv3418c), *groEL1* (Rv3417c), and *groEL2* (Rv0440); and a fourth gene, Rv0991c, is uncharacterized (36) (Fig. 2A). Chaperonins, which are represented in all domains of life, facilitate the folding of protein substrates (37, 38). In bacteria, Hsp60-type chaperonins are composed of two stacked, heptameric rings of Hsp60 (GroEL) subunits, forming a chamber in which proteins fold; the chamber is capped by a heptamer of Hsp10 (GroES) subunits [reviewed in (39)]. *M. tuberculosis* is unusual among prokaryotes by encoding two GroEL homologues, both of which form complexes that are likely capped by GroES subunits (40). It was previously shown that an *M. tuberculosis* strain with a deletion-disruption in *hrcA* exhibits high expression of *groEL1, groEL2*, and *groES* genes, the products of which are detectable in cell lysates by Coomassie brilliant blue staining of sodium dodecyl sulfate-polyacrylamide (SDS-PAGE) gels (36). Using this method, we observed protein species corresponding to GroEL1 or GroEL2 that were more abundant in an *mpa hrcA* double mutant (D*mpa::hyg hrcA::*MycoMarT7) (Fig. 2B) and in an *hrcA* single mutant (∆*hrcA*::*hyg*), as well as in the other six *mpa hrcA* suppressor mutants (*SI Appendix*, Fig. S1B). We transformed an *mpa hrcA* strain with an integrative plasmid encoding *hrcA* under the control of its native promoter, creating strain MHD1344 (*SI Appendix*, Table S1). Complementation of the *hrcA* mutation successfully restored GroEL to lower, WT levels (Fig. 2B), and reversed the suppressor phenotypes, as observed by failed growth in PB-nitrate (Fig. 2C) and excessive nitrite secretion (Fig. 2D). Collectively, these data support the hypothesis that the expression of one or more genes of the HrcA regulon is required for nitrite reduction, and that HrcA regulon expression is reduced in PPS mutants.

**Figure 2.**
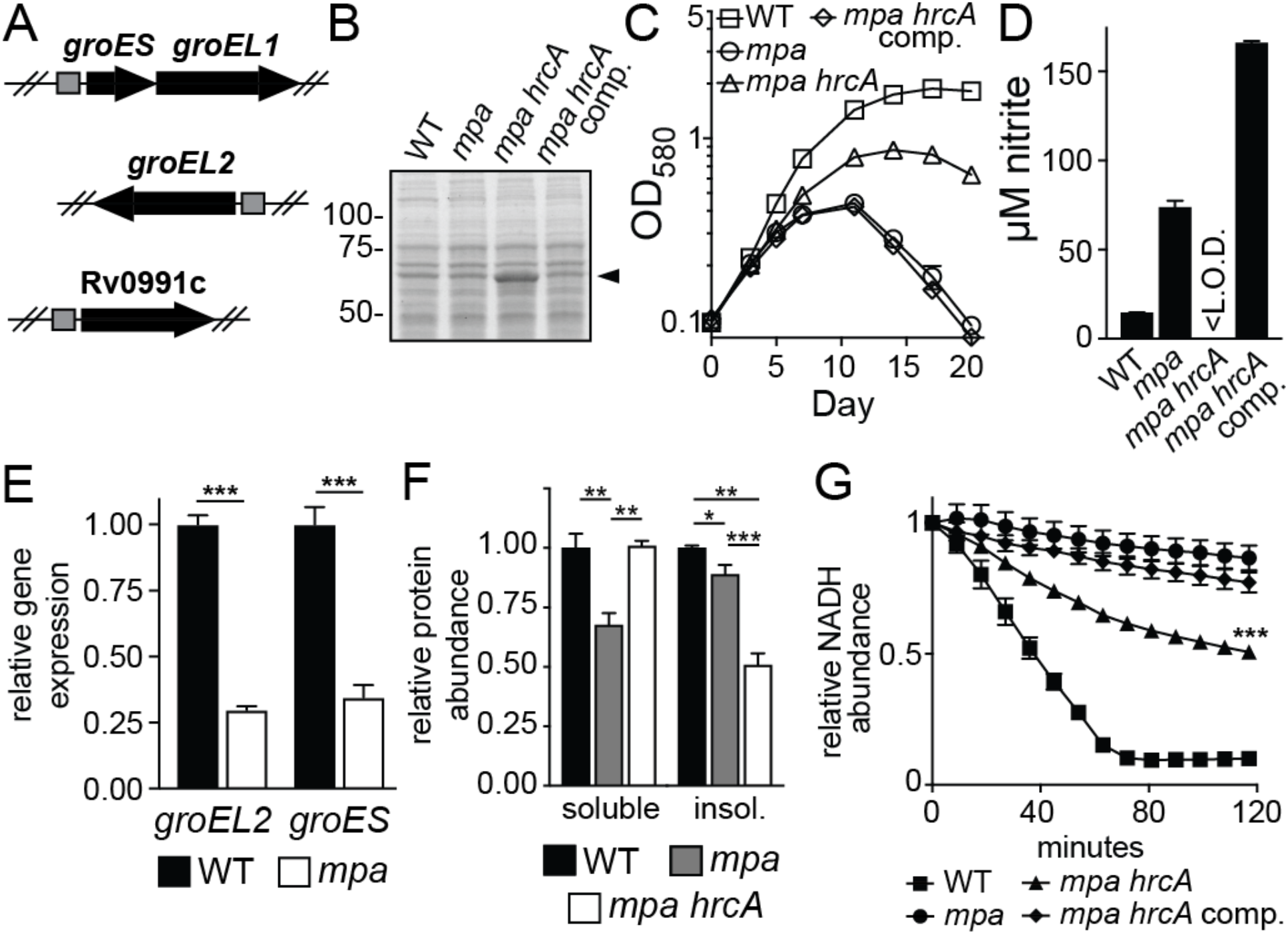
Disruption of *hrcA* increases chaperonin production and restores nitrite reductase activity to an *mpa* mutant. (A) The *M. tuberculosis* HrcA regulon, illustrating operons repressed by HrcA. Positions of HrcA consensus binding sites are shown as gray squares (36). (B) A transposon mutation in *hrcA* results in the overproduction of chaperonins. Lysates from WT (MHD1350), *mpa* (MHD1352), *mpa hrcA* (MHD1347), and *mpa hrcA* complemented (MHD1344) strains were separated by SDS-PAGE, and proteins were stained with Coomassie brilliant blue. Molecular weight markers are indicated at left. Characteristic migration pattern of GroEL1 and GroEL2, which are similar in size, is indicated with an arrowhead. (C) Disruption of *hrcA* partially rescues the growth defect of an *mpa* mutant in PB-nitrate. (D) Disruption of *hrcA* returns bacterial nitrite secretion to WT levels in an *mpa* mutant during growth in PB-nitrate. <L.O.D., below the limit of detection. (E) The chaperonin genes are transcriptionally repressed in an *mpa* mutant (MHD149) compared to the WT parental strain. (F) The *mpa* mutant contains less soluble protein than WT or *mpa hrcA* (MHD1297) strains during growth in PB-nitrate. Additionally, the *mpa hrcA* strain contains less insoluble protein than WT or *mpa* strains. (G) An *mpa* mutant is defective in nitrite reduction, which is partially rescued by *hrcA* disruption. Nitrite reductase activity was measured in normalized protein extracts from bacteria grown in PB-nitrate. Extracts were supplemented with NADH, NAD^+^, and nitrite, and nitrite reductase activity was assessed by measuring NADH oxidation over time (the nitrite reductase NirBD catalyzes electron transfer from NADH to nitrite to generate ammonium) (50). The difference in nitrite reductase activity between *mpa* and *mpa hrcA* lysates was determined to be statistically significant as indicated. Experiments in (C) through (G) each contain data from three replicate cultures. Statistical significance is indicated as follows: *, *p* < 0.05; **, *p* < 0.01; ***, *p* < 0.001. For all panels, “comp” indicated complementation with *hrcA*.

To assess gene expression in a PPS mutant, we performed a global transcriptional analysis of WT and *mpa* strains grown in PB-nitrate by RNA sequencing (RNA-Seq) (see Experimental Procedures). Because an *mpa* mutant cannot productively grow in this media (Fig. 1C), we prepared RNA from cultures grown to early logarithmic phase [optical density at 580 nm (OD_580_)=0.3]. RNA-Seq demonstrated that *groES* and *groEL2* were repressed in an *mpa* mutant compared to the parental WT strain (Fig. 2E). The remaining genes in the HrcA regulon, *groEL1* and Rv0991c, were also significantly repressed in an *mpa* mutant, although by a factor of less than two-fold (Dataset S1). This analysis suggested HrcA might be a PPS substrate.

To determine if the repression of the chaperonin genes leads to changes in protein abundance, we measured global protein levels in WT and *mpa* strains grown in PB-nitrate using tandem-mass-tag (TMT)-based quantitative mass spectrometry (MS) (see Experimental Procedures). Quantitative MS demonstrated that both GroEL2 and GroES were significantly less abundant in the *mpa* mutant compared to the WT strain; GroEL1 and Rv0991c levels were not significantly changed (Dataset S2).

*groES* and *groEL2* are essential (41), thus we were unable to disrupt these genes to test their requirement for growth in PB-nitrate. However, previous work identified a mutant with a transposon insertion in *groEL1* (1), and additionally, we deleted and disrupted Rv0991c (∆Rv0991c::*hyg*) (*SI Appendix*, Table S1). Unlike a PPS mutant, the *groEL1* and Rv0991c mutants grew well in PB-nitrate (*SI Appendix*, Fig. S2A). These data suggest that the GroES-GroEL2 ("GroESL2") complex was needed for efficient nitrite reduction.

### Chaperonin production promotes NirBD activity in *M. tuberculosis*

In order to begin to understand the association between chaperonins and nitrite reduction, we first checked if NirB or NirD abundance varied in WT, *mpa* and *mpa hrcA M. tuberculosis* strains in our MS data set (Dataset S2). While we observed a significant decrease in NirB abundance in an *mpa* mutant compared to the WT strain, this phenotype was not reversed upon disruption of *hrcA*; a similar trend was observed for NirD (*SI Appendix*, Fig. S2B). We thus concluded that changes in NirBD abundance alone could not explain the differences in nitrite reduction between the WT and *mpa* strains.

Bacterial chaperonins are required for folding many newly translated proteins, as well as for counteracting protein misfolding and aggregation under certain stress conditions (42-45). Consistent with this function of chaperonins, we always recovered less soluble protein from cell lysates of an *mpa* mutant than from the WT strain, a phenotype that was rescued by *hrcA* disruption (Fig. 2F). Based on these data, we hypothesized that chaperonins promote the activity of many *M. tuberculosis* proteins, including NirBD. To test this hypothesis, we measured NirBD activity in *M. tuberculosis* extracts. Bacterial extracts were supplemented with excess substrate (nitrite) and nicotinamide adenine dinucleotide (NAD), a cofactor that is required for NirBD activity (46). Compared to extracts made from the WT strain, nitrite reduction in *mpa* mutant extracts was at or below the limit of detection. Importantly, we observed a partial restoration of activity in *mpa hrcA* mutant extracts (Fig. 2G). This result suggested there was an intrinsic defect in NirBD activity in the *mpa* mutant that was restored by chaperonin overproduction. Notably, the incomplete rescue of nitrite reductase activity in an *mpa hrcA* strain might be explained by our observation that NirB levels were not restored by disruption of *hrcA* (*SI Appendix*, Fig. S2B).

### HrcA can be pupylated *in vitro*

The observations that the HrcA regulon was repressed in an *mpa* mutant (Fig. 2E and Dataset S1) and that the disruption of *hrcA* rescued a growth defect of a PPS mutant (Fig. 2C) suggested that HrcA might be a proteasome substrate. To test this hypothesis, we first determined if HrcA could be pupylated *in vitro*. We purified *M. tuberculosis* HrcA with carboxyl-terminal FLAG and hexahistidine (His_6_) tandem-affinity tags (HrcA_TAP_) from *Escherichia coli* (*E. coli*). Following incubation of HrcA_TAP_ with purified His_6_-Pup_Glu_ and PafA-His_6_, which are sufficient to pupylate proteins, we observed the appearance of a higher-molecular weight species corresponding to the expected size of His_6_-Pup~HrcA_TAP_ (Fig. 3A, compare lanes 1 and 2). In *M. tuberculosis*, proteins are usually pupylated at a specific lysine (16); we thus attempted to identify a pupylation site on HrcA. We made several HrcA_TAP_ variants, each with lysine-to-arginine mutations in one or two of the six lysines in HrcA. Surprisingly, no single lysine was essential for the *in vitro* pupylation of HrcA_TAP_ (Fig. 3A, lanes 3 through 7). Meanwhile, substitution of all six lysines abolished HrcA_TAP_ pupylation (Fig. 3A, lane 8). Importantly, because the epitope tag on HrcA_TAP_ contained two non-native lysines, this experiment demonstrated some substrate specificity for HrcA pupylation by PafA.

**Figure 3.**
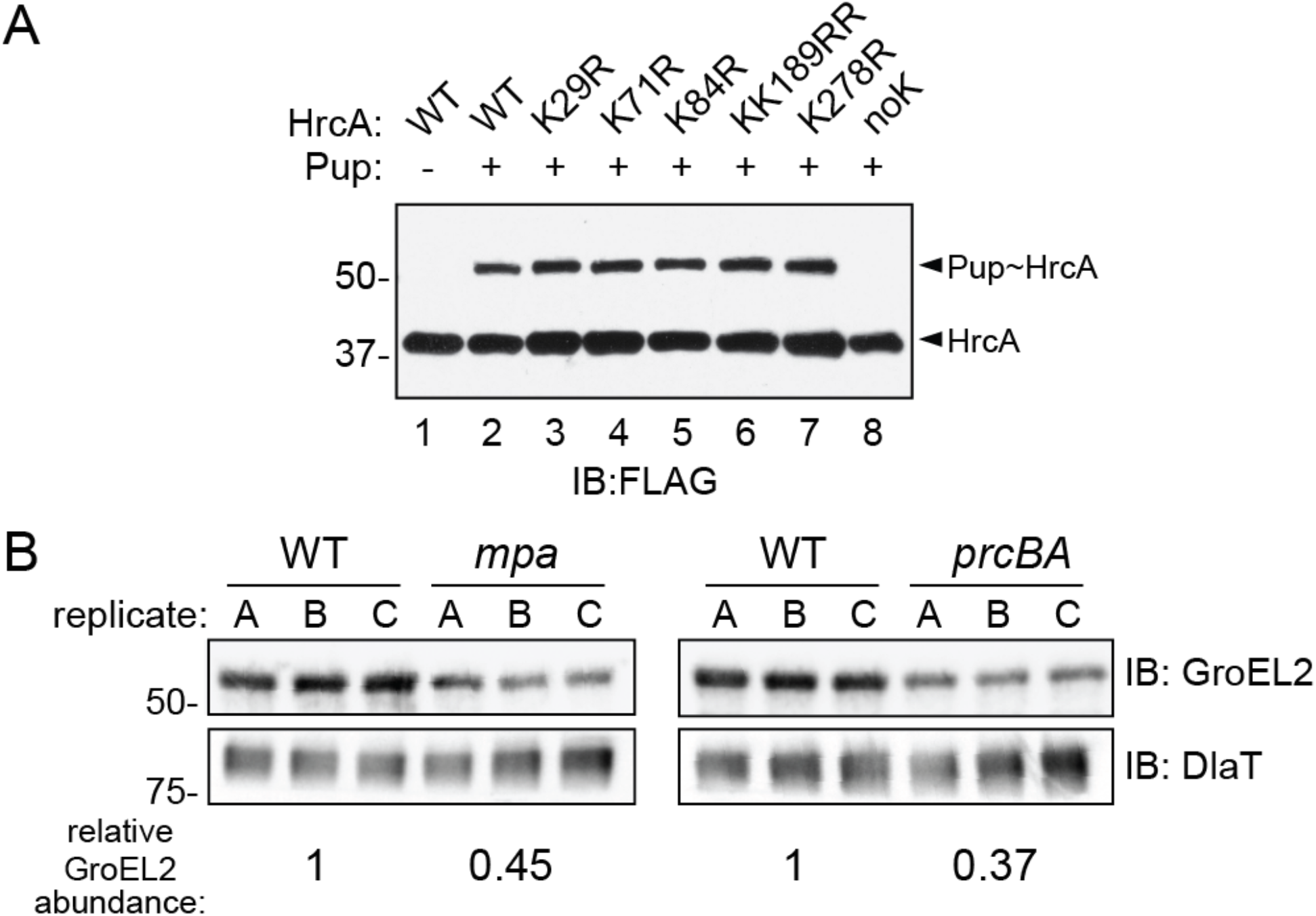
HrcA is a pupylated protein that is likely degraded by the *M. tuberculosis* proteasome. (A) Purified HrcA can be pupylated on any of its lysines by PafA. His_6_-Pup_Glu_ and PafA-His_6_ were co-incubated with HrcA_TAP_ WT or lysine-to-arginine (K>R) variants, and both native and pupylated HrcA were detected by immunoblotting (IB) using an antibody that recognizes an affinity tag on HrcA_TAP_ (FLAG). Data are representative of three independent experiments. (B) GroEL2 abundance is low in both *mpa* (MHD149) and *prcBA* strains compared to the WT parental strain. Immunoblots for GroEL2 and dihydrolipoamide acyltransferase (DlaT) were performed on the same membrane using samples obtained from replicate PB-nitrate cultures. For each lane, GroEL2 was normalized to DlaT, a protein that is not regulated by the PPS. The difference in normalized GroEL2 abundance between strains is indicated at the bottom; for comparison of WT and *mpa* strains, this difference had a *p*-value of 0.07; the difference in GroEL2 abundance between WT and *prcBA* strains had a *p*-value of < 0.01.

We sought to test if the pupylation of HrcA leads to its degradation *in vivo*. However, we were unable to observe endogenous HrcA in *M. tuberculosis* under any condition. We were unsuccessful in generating antibodies to detect HrcA in *M. tuberculosis* lysates, and HrcA was barely detected by TMT-based quantitative MS (Dataset S2). We also tried to use an epitope-tagged HrcA allele, but the tag abolished its repressor function. Finally, we introduced an *hrcA* allele lacking all of its lysines into an *hrcA* null mutant; however, this *hrcA* allele also completely lost its repressor activity.

Despite the technical limitations preventing us from observing pupylation or degradation of HrcA *in vivo*, our observation that *mpa*, *pafA* or *prcBA* mutants could not grow in nitrate (Fig. 1E) suggested that optimal chaperonin expression requires PPS-dependent proteolysis. We therefore predicted that the HrcA regulon would be repressed similarly in the *mpa* and *prcBA* mutants. We compared the abundance of GroEL2 in lysates from WT, *mpa*, and *prcBA* strains grown in PB-nitrate. We observed a similarly low abundance of GroEL2 in both the *mpa* and *prcBA* strains compared to the WT strain (Fig. 3B). Collectively, the genetic evidence along with the pupylation assays suggest that the degradation of HrcA is necessary for maintaining chaperonin gene expression in *M. tuberculosis* grown in nitrate.

### Gain-of-function mutations in *nadD* rescue a defect in NAD availability in an *mpa* mutant

We identified four different point mutations in *nadD* (nicotinate mononucleotide adenylyltransferase) (*SI Appendix*, Table S1) that each rescued the growth of an *mpa* mutant in PB-nitrate (Fig. 4A). NadD catalyzes a committed step in the biosynthesis of NAD, a molecule that serves as an electron carrier in a wide variety of essential redox reactions (47). Through ATP hydrolysis, NadD transfers adenosine monophosphate to nicotinic acid mononucleotide (NaMN), generating nicotinic acid adenine dinucleotide (NaAD), a direct precursor to NAD (48). In *M. tuberculosis*, NadD is constitutively required for the production of NAD (49).

**Figure 4.**
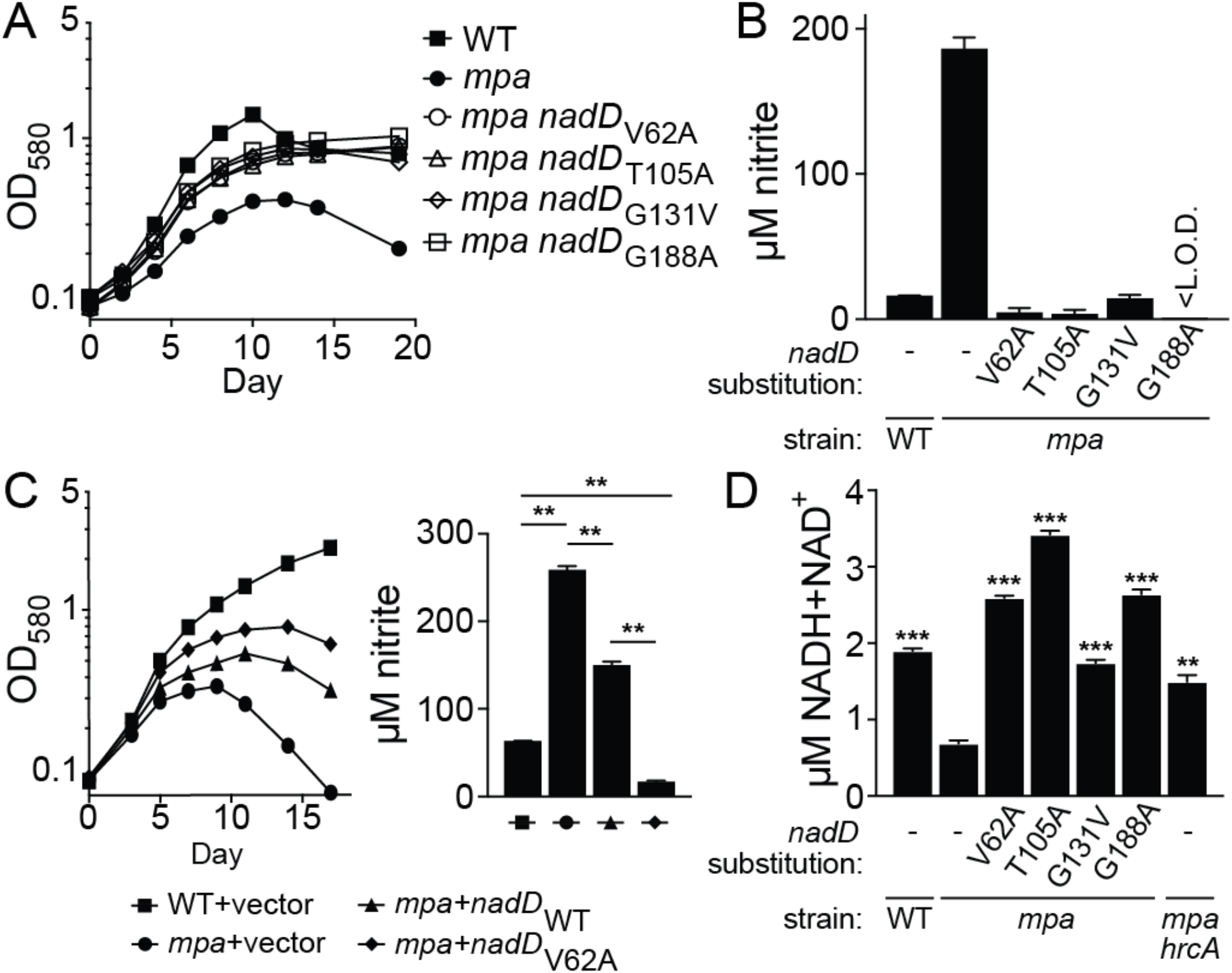
Point mutations in *nadD* restore nitrate growth to an *mpa* mutant and increase NAD abundance in bacteria. (A) Amino acid substitutions in NadD partially rescue growth of an *mpa* mutant in PB-nitrate. Strains WT, MHD149, MHD1294, MHD1300, MHD1301, and MHD1311 are represented. Note that strains MHD1294, MHD1300, MHD1301, and MHD1311 each have transposon insertions in unrelated genes (see *SI Appendix*, Table S1 for full genotypes). (B) *nadD* mutations restore nitrite secretion by the *mpa* strain to WT levels during growth in PB-nitrate. (C) Ectopic expression of WT *nadD* or *nadD*_V62A_ partially rescues growth of an *mpa* mutant in PB-nitrate (left) and lowers nitrite secretion by the *mpa* mutant (right). Strains MHD1350, MHD1352, MHD1440, and MHD1456 are represented. (D) An *mpa* mutant contains less NAD than a WT strain, a defect that is rescued both by mutations in *nadD* and by disruption of *hrcA* (MHD1297). Total NAD [oxidized (NADH) and reduced (NAD^+^) forms] was quantified in lysates of bacteria grown in PB-nitrate; statistical significance is indicated by comparison to the *mpa* single mutant. Experiments in (A) through (D) each contain data from three replicate cultures. **, *p* < 0.01; ***, *p* < 0.001.

The four mutations we identified in *nadD* resulted in the amino acid substitutions V62A, T105I, G131V, and G188A (V, valine; A, alanine; T, threonine; I, isoleucine; G, glycine); two additional strains, recovered from independent mutant pools, also encoded a NadD_V62A_ allele (see *SI Appendix*, Table S1). Consistent with their ability to grow in PB-nitrate, all four *mpa nadD* suppressor strains secreted low nitrite levels comparable to the parental WT strain, demonstrating that NirBD activity was restored in these strains (Fig. 4B).

Because NadD is essential for the growth of *M. tuberculosis* (49), we predicted that these *nadD* suppressor mutations resulted in a gain of function. To test this hypothesis, we transformed a single copy of either WT *nadD* or *nadD*_V62A_ into an *mpa* strain and assessed growth of these transformants in PB-nitrate. As expected, ectopic expression of *nadD*_V62A_ partially rescued the growth of the *mpa* parental strain, while ectopic expression of WT *nadD* had an intermediate phenotype. Likewise, ectopic expression of *nadD*_V62A_ had a dominant effect to reduce nitrite secretion, even in the presence of the endogenous, WT *nadD* (Fig. 4C). We also measured the levels of total oxidized and reduced NAD (NAD^+^ and NADH, respectively) in *M. tuberculosis* lysates from our strains. Interestingly, we observed a nearly three-fold reduction in NAD abundance in an *mpa* mutant relative to the parental WT strain. Importantly, all four *nadD* mutations restored NAD abundance in an *mpa* mutant to levels equal to or greater than that of the WT strain (Fig. 4D). Thus, *nadD* gain-of-function mutations rescued a defect in NAD availability in the *mpa* strain.

NAD depletion could affect many redox-associated enzymes in *M. tuberculosis*; however, there exists a direct link between NAD and nitrate catabolism. NirBD catalyzes electron transfer from NADH to nitrite, producing ammonium and NAD^+^ (50). This reaction also requires the presence of NAD^+^ itself (46). Accordingly, the reduced levels of NAD in the *mpa* mutant could contribute to this strain’s defect in NirBD activity.

We sought to understand the molecular basis by which amino acid substitutions in NadD result in increased production of NAD *in vivo*. We produced WT and variant (V62A, T105I, G131V, and G188A) NadD in *E. coli*, and purified these proteins to homogeneity. We first measured NadD protein stability using a thermal shift assay (see Experimental Procedures). Remarkably, three of the four NadD variants (T105I, G131V, G188A) displayed a more than 13ºC increase in melting temperature compared to WT NadD (Table 1 and *SI Appendix*, Fig. S3A). We next measured the rate of ATP hydrolysis by WT and variant NadD upon incubation with NaMN. While NadD G131V and G188A hydrolyzed ATP at a higher rate, the remaining two NadD variants had less activity than the WT enzyme (Table 1).

**Table 1.** Analysis of NadD variants.

| Protein Sample | T<sub>m</sub> (s.d.)<sup>a</sup>, °C | Activity (s.d.), nmol min<sup>-1</sup> mg<sup>-1</sup> |
| --- | --- | --- |
| NadD WT | 54.4 (1.06) | 1.09 (0.01) |
| NadD V62A | 48.7 (0.94) | 0.48 (0.015) |
| NadD T105I | 68.0 (1.21) | 0.11 (0.05) |
| NadD G131V | 68.9 (0.82) | 1.95 (0.005) |
| NadD G188A | 67.7 (0.78) | 2.34 (0.07) |

<sup>a</sup>T<sub>m</sub>, melting temperature; s.d., standard deviation.

We mapped the amino acid substitutions on the NadD crystal structure. T105I and G131V are located in the core of the NadD monomer (*SI Appendix*, Fig. S3B). This region is characterized by hydrophobic interactions between a central β-sheet and several α-helices (51); accordingly, such hydrophobic amino acid substitutions may stabilize NadD by increasing core packing, which might explain their increased thermal stability. In contrast, substitutions V62A and G188A lie at a subunit-to-subunit interface in the NadD tetramer (*SI Appendix*, Fig. S3B). NadD forms both dimers and tetramers *in vitro* (51); while the state of NadD assembly *in vivo* is unknown, it is possible that substitutions at the surface of NadD monomers influence the oligomeric state of NadD to affect its catalytic activity (either positively or negatively) in *M. tuberculosis*. While we cannot yet explain why two of the NadD mutant alleles show slower activity *in vitro*, our genetic data suggest NadD activity is higher *in vivo* for all four mutants.

We also found that the low NAD levels in an *mpa* mutant were also restored by disruption of *hrcA* (Fig. 4D). It is possible that either the HrcA regulon is needed to support NAD synthesis, or that in the absence of Mpa function, one or more NAD-consuming enzymes deplete the cellular stores of this cofactor.

### Nitrogen sources affect steady state pupylome levels

Our results up to now suggest that the PPS degrades HrcA to allow for the expression of chaperonin genes in bacteria growing in nitrate. However, we did not know whether these observations reflected the specific degradation of HrcA, or a mass degradation of substrates by the proteasome. To address this question, we grew *M. tuberculosis* in PB-Asn, which permits robust growth of an *mpa* mutant (Fig. 1B), or in PB-nitrate and quantified the abundance of pupylated proteins in bacterial lysates detectable by immunoblotting. We observed a nearly two-fold decrease in pupylome abundance in bacteria grown in PB-nitrate compared to PB-Asn. We also observed a decrease in the abundance of (unpupylated) inositol-3-phosphate synthase (Ino1), a model PPS substrate (16), but not of PrcB (Fig. 5A). This result suggested that there was an increase in the degradation of pupylated proteins, rather than a decrease in pupylation, during growth in PB-nitrate. To further test this point, we used a reporter protein, Pup-Zur-His_6_, to specifically observe the degradation of a “prepupylated” protein in *M. tuberculosis*. Zur (zinc uptake regulator, Rv2359) is an *M. tuberculosis* protein that lacks lysines, and therefore cannot be pupylated *in vivo*. Pup is translationally fused to Zur through a linear amide, rather than an isopeptide bond, and cannot be depupylated; thus, the abundance of this reporter specifically assesses proteolysis by the Mpa-proteasome (52). In WT *M. tuberculosis*, we observed a decrease in Pup-Zur-His_6_ abundance in bacteria grown in PB-nitrate compared to bacteria cultured in PB-Asn. Meanwhile, in an *mpa* mutant, there was no difference in Pup-Zur-His_6_ levels upon growth in either medium, supporting a model whereby the Mpa-proteasome degrades pupylated proteins during growth in nitrate (Fig. 5B).

**Figure 5.**
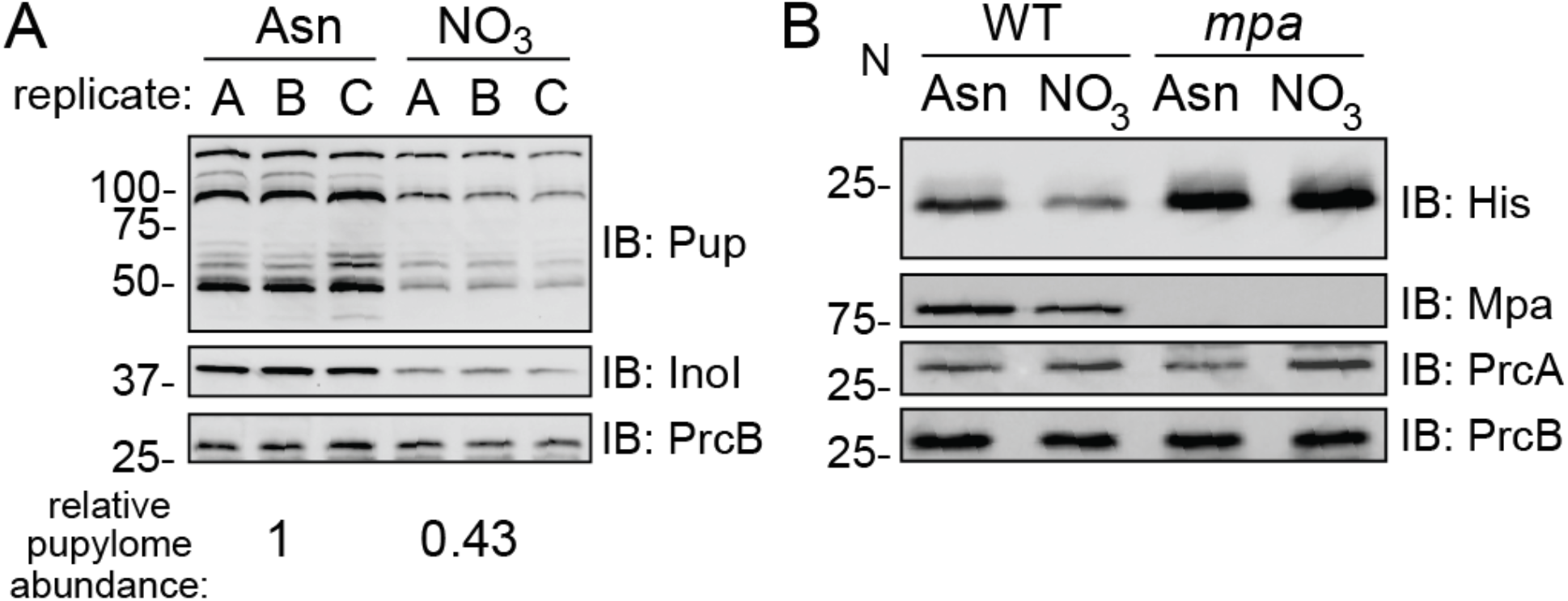
Abundance of pupylated proteins depends on the nitrogen source. (A) *M. tuberculosis* contains a lower abundance of pupylated proteins, and of a model PPS substrate, when grown in PB-nitrate compared to PB-Asn. Pupylated proteins were detected by immunoblot (IB) using a monoclonal antibody that recognizes *M. tuberculosis* Pup. The same immunoblot membranes were used to detect inositol-3-phosphate synthase (Ino1) and PrcB. The relative pupylome abundance between growth conditions (bottom) was normalized by PrcB abundance and was statistically significant (*p* <0.05). (B) Pup-Zur-His_6_ levels are reduced in WT *M. tuberculosis* grown in PB-nitrate compared to PB-Asn. The normalized Pup-Zur-His_6_ intensity in the second lane relative to the first lane is 0.62. PrcB and Mpa were detected on the same membrane, while PrcA was detected using a membrane separately prepared with the same lysates.

Previous work has shown that total nitrogen starvation is associated with a decrease in pupylated proteins in *M. smegmatis*. This phenomenon was attributed to a greater abundance of 20S CPs upon nitrogen starvation, suggesting that *M. smegmatis* regulates the production of the degradation machinery in response to nitrogen availability (26). However, we found that while the abundance of the pupylome, Ino1, and Pup-Zur-His_6_ decreased during growth in PB-nitrate, the levels of Mpa, PrcA, and PrcB remained unchanged (Fig. 5B). Therefore, we propose that the regulation of proteasomal degradation in *M. tuberculosis* during growth in nitrate does not require significantly altering levels of the known proteolytic components.

## DISCUSSION

In this work, we established that *M. tuberculosis* requires an intact PPS to assimilate nitrogen from nitrate. Specifically, the ability of *M. tuberculosis* to reduce nitrite depended on the expression of the Hsp60 chaperonin genes, including *groES* and *groEL2*. We found that HrcA, a repressor of the *groES and groEL1/2* genes, is most likely pupylated and degraded by the proteasome in order to allow for the production of the GroESL2 complex during growth in nitrate. Additionally, we found that NAD levels were reduced in the absence of a functional PPS, which could also contribute to the observed defect in nitrite reduction in PPS mutants. Lastly, we showed that the abundance of PPS substrates changed depending on the nitrogen source provided to *M. tuberculosis*.

While mouse models of *M. tuberculosis* infection have demonstrated a requirement for bacterial uptake of asparagine and aspartate as nitrogen sources (30, 31), the importance of nitrate as a nutrient during infection is less clear. An *M. tuberculosis narG* mutant, which is unable to reduce nitrate, is fully virulent in mice (53). However, unlike in humans, *M. tuberculosis* lesions in most inbred mouse lines are not hypoxic (53, 54); because nitrate import and reduction occur most abundantly under anaerobic conditions (55-57), these infection models may not accurately reflect nitrate utilization during a human infection.

Our results suggest that *M. tuberculosis* NirBD activity requires de-repression of the HrcA regulon and support a model by which HrcA is degraded in a PPS-dependent manner (Fig. 6). Studies of HrcA from other bacterial species have shown that this repressor acts as a thermosensor: an increase in temperature induces the dissociation of HrcA from DNA, presumably allowing for the expression of factors necessary to respond to heat-induced protein misfolding (58, 59). We observed that the PPS alleviates HrcA repression in the absence of heat shock, suggesting that there are other ways of inducing the expression of the *hsp60* protein quality control genes in *M. tuberculosis*. Importantly, it is unknown how many *M. tuberculosis* proteins depend on GroESL2 for folding. The identification of other GroESL2 substrates could potentially uncover additional pathways whose function depends on PPS-mediated control of chaperonin gene expression. Importantly, these pathways may at least partially explain how defects in the PPS lead to highly attenuated bacteria in animals.

**Figure 6.**
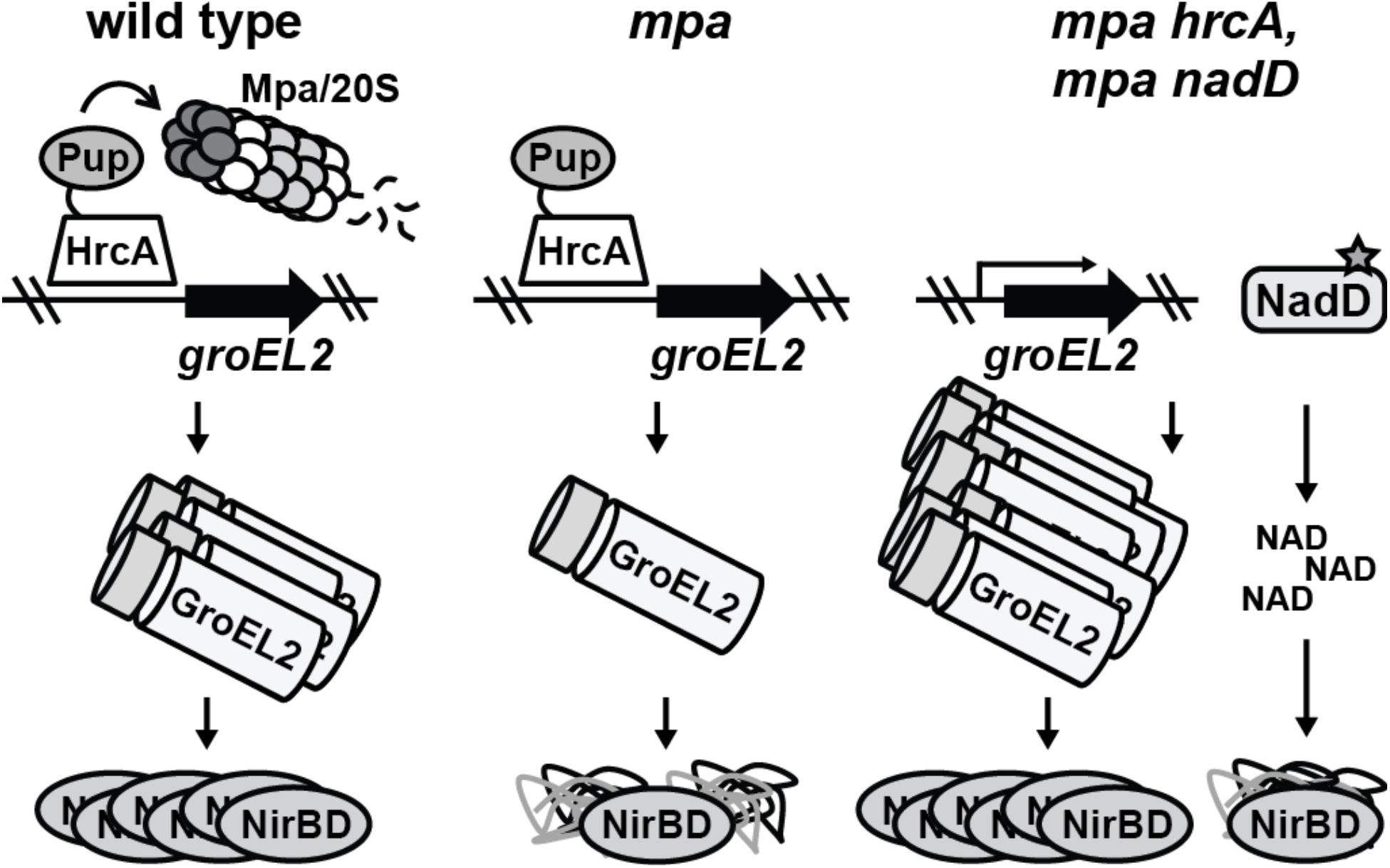
Model of PPS control over *M. tuberculosis* nitrate metabolism. Left: HrcA, which represses the *M. tuberculosis* chaperonin genes including *groEL2*, is likely pupylated and degraded by the Mpa/20S CP proteasome to allow for the full expression of the chaperonins that promote the folding or assembly of the nitrite reductase NirBD. Middle: Failure of proteasomal degradation in *M. tuberculosis* leads to the repression of the chaperonin genes, preventing the formation of functional NirBD. Right: Disruption of *hrcA* restores NirBD activity in an *mpa* mutant through the full de-repression the chaperonin genes, while gain-of-function mutations in *nadD* increase the abundance of NAD to promote NirBD catalysis.

According to the most well-characterized model of chaperonin activity in *E. coli*, misfolded or unfolded proteins become encapsulated within a GroES-GroEL chamber, a hydrophobic space in which substrates fold (37, 38). NirD has a mass of 12.5 kD, a size that is within the range of most *E. coli* GroEL substrates (60); in contrast, the 90 kD NirB subunit is too large to be fully encapsulated. Nonetheless, a mechanism of chaperonin-mediated folding of large proteins without encapsulation has been described (61, 62), and several high-molecular weight *E. coli* proteins have been identified as *in vivo* GroEL substrates (60). Thus, it is possible that NirB and/or NirD are endogenous substrates of GroESL2 in *M. tuberculosis*.

While a failure of *M. tuberculosis* to maintain chaperonin levels is associated with a loss of NirBD function, we have also shown that an *mpa* mutant grown in nitrate lacks WT levels of NAD, which is required for NirBD activity. Proteomic analysis of WT and *mpa* strains did not identify alterations in the abundance of any enzymes within the NAD biosynthetic pathway that could explain the failure of an *mpa* mutant to maintain WT NAD levels (Dataset S2). It is possible that the NAD pool is exhausted by the accumulation of one or more PPS substrates that consume NAD. However, it is telling that in addition to compensatory mutations in *nadD*, *hrcA* disruption is sufficient to restore NAD levels in an *mpa* mutant. Taken together, these observations suggest that GroESL2 may also promote the folding or assembly of NadD or other enzymes in the NAD synthetic pathway. In turn, the gain-of-function mutations in *nadD* that were selected for in our suppressor screen may allow NadD to remain functional in spite of reduced chaperonin levels. Importantly, NadD is thought to be essential in most bacteria, and is of interest as a drug target in a potentially diverse set of pathogens, including *M. tuberculosis* (49, 63). Thus, the novel NadD variants that we describe here highlight the importance of NadD activity under stress conditions and strengthen the potential of NadD as an underappreciated drug target.

In a study using *M. smegmatis*, Gur and colleagues found that the pupylome is less abundant during nitrogen starvation, an observation that is similar to what we observed with *M. tuberculosis* grown in nitrate broth. It was proposed that altered levels of components of the PPS are responsible for this phenotype in *M. smegmatis* (26). In contrast, we did not observe conspicuous changes in proteasome component abundance in *M. tuberculosis* despite a dramatic change in substrate abundance, suggesting that there are differences in the regulation of proteasomal activity between these bacterial species. Instead of altering PPS component levels, it is conceivable that there are post-translational modifications on the proteasome itself that alter its activity. For example, *M. tuberculosis* kinases PknA and PknB can phosphorylate PrcA and PrcB (64); although there is no indication that this activity occurs in a physiological setting, phosphorylation could potentially affect the activity of 20S CPs. Additionally, there may be factors that modulate the association of Mpa with the 20S CP, since attempts to observe a robust Mpa-20S CP interaction *in vitro* have been unsuccessful (65).

In addition to identifying a novel role of the *M. tuberculosis* PPS in regulating chaperonin and NAD levels during growth in nitrate, we found a specific condition during which proteasomal degradation appears to be stimulated. Because the Hsp60 system is undoubtedly required for the function of numerous proteins, it seems likely that other environmental cues could activate proteasomal degradation to induce *hsp60* regulon expression. Thus, the molecular mechanisms by which PPS function might be altered, as well as other growth conditions that promote proteolysis, warrant further investigation.

## EXPERIMENTAL PROCEDURES

### Bacterial strains, plasmids, primers, and culture conditions

Bacterial strains, plasmids, and primers used in this study are listed in *SI Appendix*, Table S1. Chemicals used for making all buffers and bacterial media were purchased from ThermoFisher, Inc. unless otherwise indicated. *M. tuberculosis* was grown in "7H9" [BD Difco Middlebrook 7H9 broth with 0.2% glycerol and supplemented with 0.5% bovine serum albumin (BSA), 0.2% dextrose, 0.085% sodium chloride, and 0.05% Tween-80]. For culturing *M. tuberculosis* in single nitrogen sources, a base of Proskauer-Beck (PB) minimal medium (66) with no nitrogen source ("PB-base") was prepared with 0.5% potassium phosphate monobasic, 0.06% magnesium sulfate heptahydrate, 1.5% glycerol, 0.25% magnesium citrate dibasic anhydrous, and 0.05% Tween-80. The following nitrogen sources were added to a final concentration of 10 mM: asparagine (PB-Asn), glutamate (PB-Glu), arginine (PB-Arg), sodium nitrate (PB-nitrate), or ammonium chloride (PB-ammonium); pH was adjusted to 6.4 after nitrogen addition. PB broths were autoclaved or filtered prior to use. *M. tuberculosis* was incubated at 37°C for all experiments.

For solid media, *M. tuberculosis* was grown on "7H11" agar (BD Difco Middlebrook 7H11) containing 0.5% glycerol and supplemented with 10% final volume of BBL Middlebrook OADC Enrichment. For selection of *M. tuberculosis*, the following antibiotics were used as needed: kanamycin 50 µg/ml, hygromycin 50 µg/ml, and gentamicin 15 µg/ml.

*E. coli* was cultured in BD Difco Luria-Bertani (LB) broth or on LB-Agar. Media were supplemented with the following antibiotics as needed: kanamycin 100 µg/ml, hygromycin 150 µg/ml, and gentamicin 15 µg/ml.

For all experiments in which *M. tuberculosis* was cultured in PB broth, bacteria were first grown in 7H9 to an OD_580_ of 0.5 - 0.8, washed three times in PBS-T [phosphate buffered saline (Corning) with 0.05% Tween-80], and resuspended in the appropriate PB broth. For growth curves, bacteria were harvested by centrifugation at 500 × *g* for five minutes to remove large clumps of bacteria prior to dilution into fresh broth.

A detailed description of plasmid construction is provided in *SI Appendix*, Supplementary Experimental Procedures.

### Protein purification, antibody production, and immunoblotting

Purification of PafA-His_6_ and His_6_-Pup_Glu_ was described previously (9, 15). HrcA was made with a C-terminal affinity tag consisting of FLAG and His_6_ epitopes separated by a five-amino acid linker ("HrcA_TAP_"). *M. smegmatis* PrcB was made with a C-terminal His_6_ (smPrcB-His_6_). HrcA_TAP_, smPrcB-His_6_, and PrcA-His_6_ were produced in *E. coli* strain ER2566 and purified by affinity chromatography using Ni-NTA agarose (Qiagen) according to the manufacturer’s instructions (PrcA and PrcB were purified under urea denaturing conditions). To make rabbit polyclonal immune serum, approximately 200 µg PrcA-His_6_ or smPrcB was used to immunize rabbits (Covance, Denver, PA). Purification of recombinant NadD is described in *SI Appendix*, Supplementary Experimental Procedures. Antibodies to *M. tuberculosis* DlaT were a gift from R. Bryk and C. Nathan.

Separation of proteins in *in vitro* assays and in *M. tuberculosis* lysates was performed using 10% sodium dodecyl sulfate-polyacrylamide (SDS-PAGE) gels, with the exception of the experiment shown in Fig. 5B, which used a 15% SDS-PAGE gel. Bio-Safe Coomassie Stain (Bio-Rad) was used to stain gels. For immunoblots, proteins were transferred from SDS-PAGE gels to nitrocellulose membranes (GE Amersham), and analyzed by immunoblotting as indicated. Detailed immunoblotting procedures are found in *SI Appendix*, Supplementary Experimental Procedures.

To quantify GroEL2 abundance in Fig. 3B, we used ImageJ (https://imagej.nih.gov) to measure the pixel density of GroEL2 and DlaT signals in immunoblot images. To normalize each lane, the GroEL2 density was divided by the DlaT density. Quantification of pupylome and Pup-Zur-His_6_ abundances in Fig. 5 was performed in the same manner, using the PrcB signal to normalize the pupylome or His signal for each lane. For the fractionation experiment shown in Fig. 2F, total protein content in soluble and insoluble lysate fractions was determined by separating samples on SDS-PAGE gels, staining gels with Coomassie brilliant blue, and using ImageJ to measure pixel density in scanned images.

### Preparation of *M. tuberculosis* extracts

To generate protein extracts for gel separation and immunoblotting, *M. tuberculosis* cultures were grown to an OD_580_ of 0.3. Equal amounts of bacteria were harvested by centrifugation, resuspended in lysis buffer (50 mM Tris, 150 mM sodium chloride, and 1 mM ethylenediaminetetraacetic acid (EDTA), pH 8.0). and transferred to a tube containing 250 µl of 0.1 mm zirconia beads (BioSpec Products). Bacteria were lysed using a mechanical bead-beater (BioSpec Products). Whole lysates were mixed with 4 × SDS sample buffer (250 mM Tris pH 6.8, 2% SDS, 20% 2-mercaptoethanol, 40% glycerol, 1% bromophenol blue) to a 1 × final concentration, and samples were boiled for 5 minutes. For preparing lysates from *M. tuberculosis* grown in 7H9, which contains BSA, an additional wash step with PBS-T was done prior to resuspension of bacteria in lysis buffer.

For the fractionation experiment shown in Fig. 2F, bacteria were lysed as described above; whole lysate was centrifuged at 16,000 × *g* for five minutes to pellet insoluble material. Supernatants were mixed with 4 × SDS sample buffer, and pellets were resuspended in 250 µl of fresh lysis buffer and mixed with 4 × SDS sample buffer; each sample was boiled for 5 minutes.

### Sequencing of suppressor mutants

*M. tuberculosis* chromosomal DNA was purified as described previously (67). Transposon insertion sites were cloned from *M. tuberculosis* genomic DNA, transformed into S17-λpir, and sequenced as previously described (1). For strains MHD149, MHD1294, MHD1300, MHD1301, MHD1302, MHD1304, MHD1306, MHD1307, MHD1308, and MHD1311, whole-genome sequencing was done by the Genome Technology Center at NYU Langone Health using an Illumina Hi-Seq platform. Reads were mapped to the H37Rv reference genome (NCBI) using BWA (http://bio-bwa.sourceforge.net/) and SAMtools (http://samtools.sourceforge.net/). Identification of nucleotide mutations was performed using HaplotypeCaller (Broad Institute).

### Quantification of nitrite reductase activity

All experiments were performed using cultures growing in PB-nitrate to an OD_580_ of 0.3. The concentration of nitrite in *M. tuberculosis* culture supernatants was measured using the Griess assay (68) by mixing supernatant 1:1 with Griess reagent (2.5% phosphoric acid, 0.5% sulfanilamide, 0.05% N-(1-napthyl)-ethylenediamine), incubating for 10 minutes at 25°C, and measuring absorbance at 550 nm (A_550_). A set of sodium nitrite solutions was used to make a standard curve for A_550_ measurements.

Direct measurement of nitrite reductase activity in *M. tuberculosis* extracts was performed as described previously, using NADH oxidation as a measure of NirBD activity (46). To eliminate background oxidation by NADH dehydrogenase, a membrane-associated complex, bacterial lysates were filtered and subjected to ultracentrifugation at 150,000 × *g* for 2 hours to remove insoluble material. Extracts were then normalized by total protein content after measuring the protein concentration using the Bio-Rad Protein Assay. NADH was measured in the reactions by A_340_. Background NADH oxidation by extracts in the absence of sodium nitrite was measured and determined to be negligible.

### Transcriptional analysis

To analyze gene expression, RNA was purified as previously described (69) from *M*. tuberculosis cultures grown in PB-nitrate to an OD_580_ of 0.3. Library preparation, sequencing by Illumina HiSeq, and analysis were performed by GENEWIZ, Inc. Sequence reads were mapped to the H37Rv genome using Bowtie2 (http://bowtie-bio.sourceforge.net/bowtie2/index.shtml). Unique gene hit counts were calculated using Subread (http://subread.sourceforge.net/), and differential gene expression analysis was performed using DeSeq2 (https://bioconductor.org/packages/release/bioc/html/DESeq2.html). To compare gene expression between strains, the Wald test was used to generate *p*-values and log_2_ fold changes. Genes with an adjusted *p*-value < 0.05 and absolute log_2_ fold change > 1 were called as differentially expressed genes for each comparison. Global gene expression analyses in WT and MHD149 strains is provided in Dataset S1. Raw sequencing data files are available in a PATRIC public workspace (https://patricbrc.org/workspace/public/shb360@patricbrc.org/shb2018).

### Mass spectrometry

For analysis of protein content in *M. tuberculosis* strains, bacteria were grown in PB-nitrate to an OD_580_ of 0.3. Equal amounts of bacteria were harvested by centrifugation, resuspended in freshly-prepared denaturing lysis buffer (100 mM Tris, 1 mM EDTA, 8M urea, pH 8.0), and lysed by bead beating. Whole lysates were centrifuged at 16,000 × g for 5 minutes to pellet the urea-insoluble material. Supernatants were centrifuged through a 0.22 µm Spin-X cellulose-acetate filter (Corning) and stored at −80ºC. Detailed methods for TMT-based quantitative MS are in *SI Appendix*, Supplementary Experimental Procedures. A comparison of global protein abundances between WT, MHD149, and MHD1297 is provided in Dataset S2. Raw peptide data is available in a PATRIC public workspace (https://patricbrc.org/workspace/public/shb360@patricbrc.org/shb2018).

### *In vitro* pupylation of HrcA

Pupylation assays were performed as described previously (10, 15). Briefly, reaction mixtures contained 1 µM His_6_-Pup_Glu_, 1 µM HrcA_TAP_, and 0.5 µM PafA-His_6_ at pH 8.0 in the presence of 5 mM ATP, 50 mM Tris, 20 mM magnesium chloride, 10% glycerol, 1 mM dithiothreitol, and 150 mM sodium chloride. Reactions were incubated overnight at 25°C.

### Quantification of NAD

Total NAD in *M. tuberculosis* lysates was quantified using the NAD/NADH Quantitation Kit (Sigma-Aldrich). Preparation of protein-free bacterial extracts and NAD quantification were performed according to the manufacturer’s instructions.

### NadD kinetics and stability assays

Thermal stability of NadD variants was measured using differential scanning fluorimetry (thermal shift assay). Differential scanning fluorimetry was performed using a CFX96 Touch Real-Time PCR detection system and the florescent dye SYPRO Orange stock concentration at a final concentration of 2 × in 96-well PCR plates. The initial fluorescence signal was measured after five minutes of temperature equilibration at 25°C followed by measurements at every 1°C min^−^1 until reaching 95°C. The wavelength of excitation and emission were 490 nm and 580 nm, respectively. For each experiment, the protein was run alone and in the presence of 10 mM Mg-ATP. Experiments were carried out with at least three samples per condition; results were expressed as mean values ± the standard error of the mean. Melting temperatures were calculated using CFX Manager 3.1 software’s d(RFU)/dT peak finder.

Reaction mixtures for the assay of nicotinic acid adenylyl transferase activity of NadD contained 100 mM HEPES-NaOH, pH 7.4, 10 mM magnesium chloride, 1 mM NaMN, 0.1 mM ATP (Sigma), 5 mU inorganic pyrophosphatase (Sigma) and 20 µg purified NadD in a total volume of 0.1 ml. Reactions were performed in a clear, flat-bottomed 96-well plate at room temperature. After incubation for 10 min., inorganic phosphate was detected using the Malachite Green assay (70).

### Statistical significance

With the exception of TMT-based proteomics and RNA-seq analyses, all *p-*values were calculated using Welch’s T-test.

## AUTHOR CONTRIBUTIONS

S.H.B., J.B.J., A.D., K.V.K., B.U., and K.H.D. designed research; S.H.B., J.B.J., A.D., C.T.C., and J.S.R. performed research; S.H.B., K.V.K., and K.H.D. wrote the manuscript.

## ACKNOWLEDGEMENTS

This work was supported by National Institutes of Health (NIH) grants R01 HL092774 and AI088075 awarded to K.H.D. S.H.B. and J.B.J. were supported by NIH grant T32 AT007180. S.H.B. also received support from the Jan T. Vilcek Endowed Fellowship Fund. K.V.K. was supported by NIH grant R03AI117361. We thank A. Osterman for providing the NadD expression construct. We thank C. Kenner, J. Li and R. Reed for assistance with NadD purifications. We thank S. Ehrt for the *prcBA* mutant and R. Copin for assistance in analysis of whole-genome sequencing data. We thank M. Samanovic, S. Zhang, A. Darwin, and members of the A. Darwin lab for helpful discussions.

## Supplementary Information for

**This PDF file includes:**

Table S1

Figures S1 to S3

Legends for Figures S1 to S3

Supplementary Experimental Procedures

References for SI Appendix

**Other supplementary materials for this manuscript include the following:**

Datasets S1 and S2

### SUPPLEMENTARY TABLES AND FIGURES

**TABLE S1.**
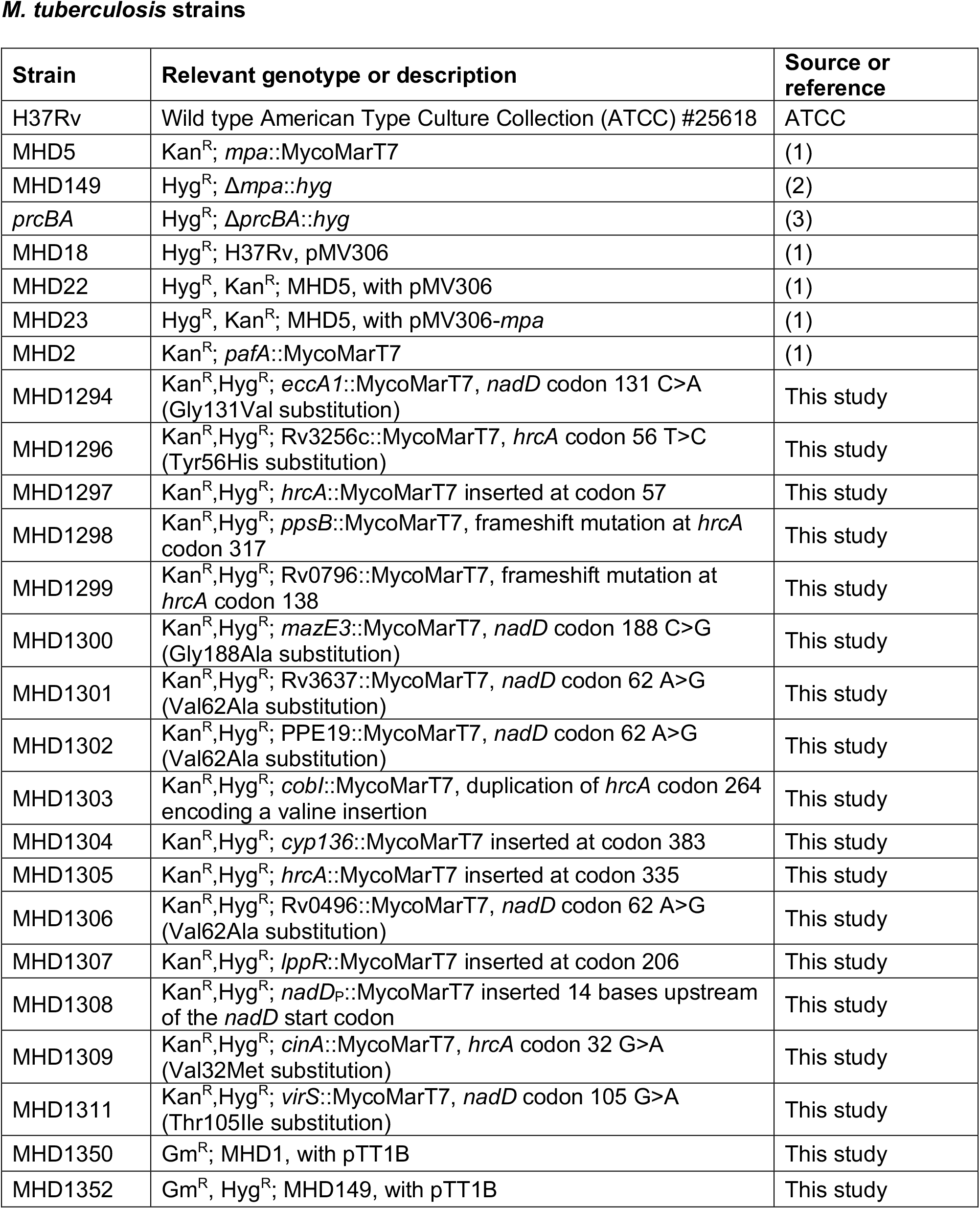
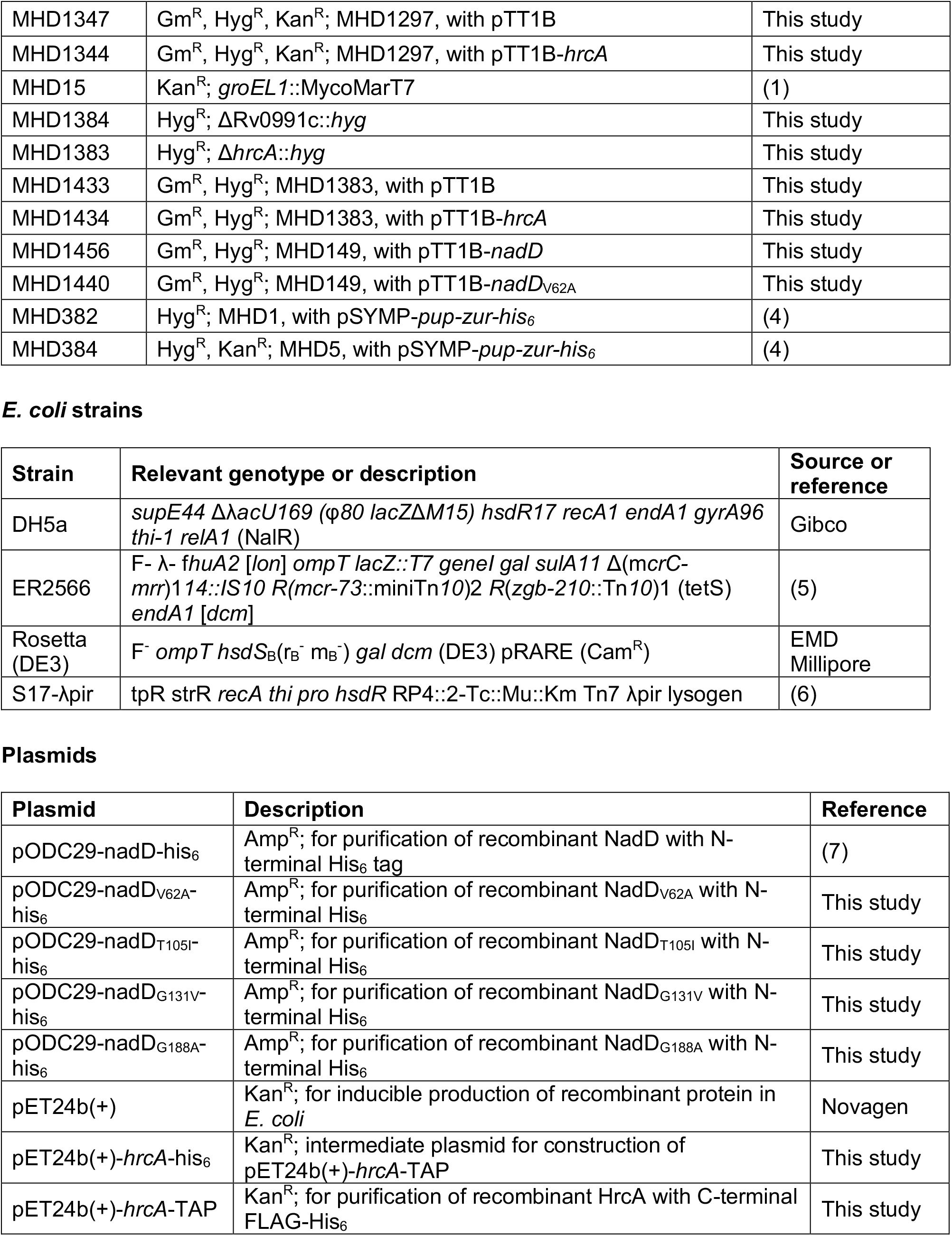
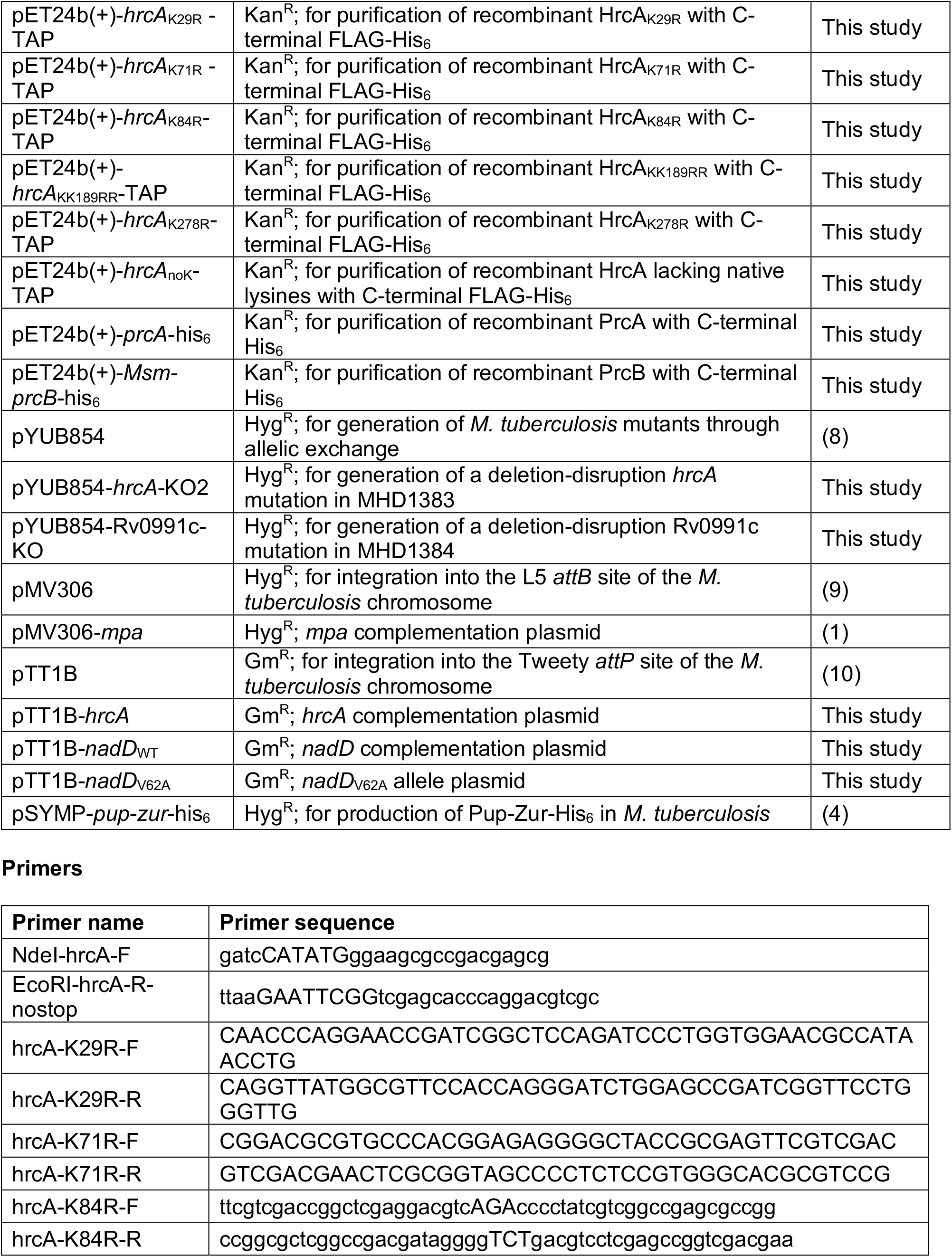
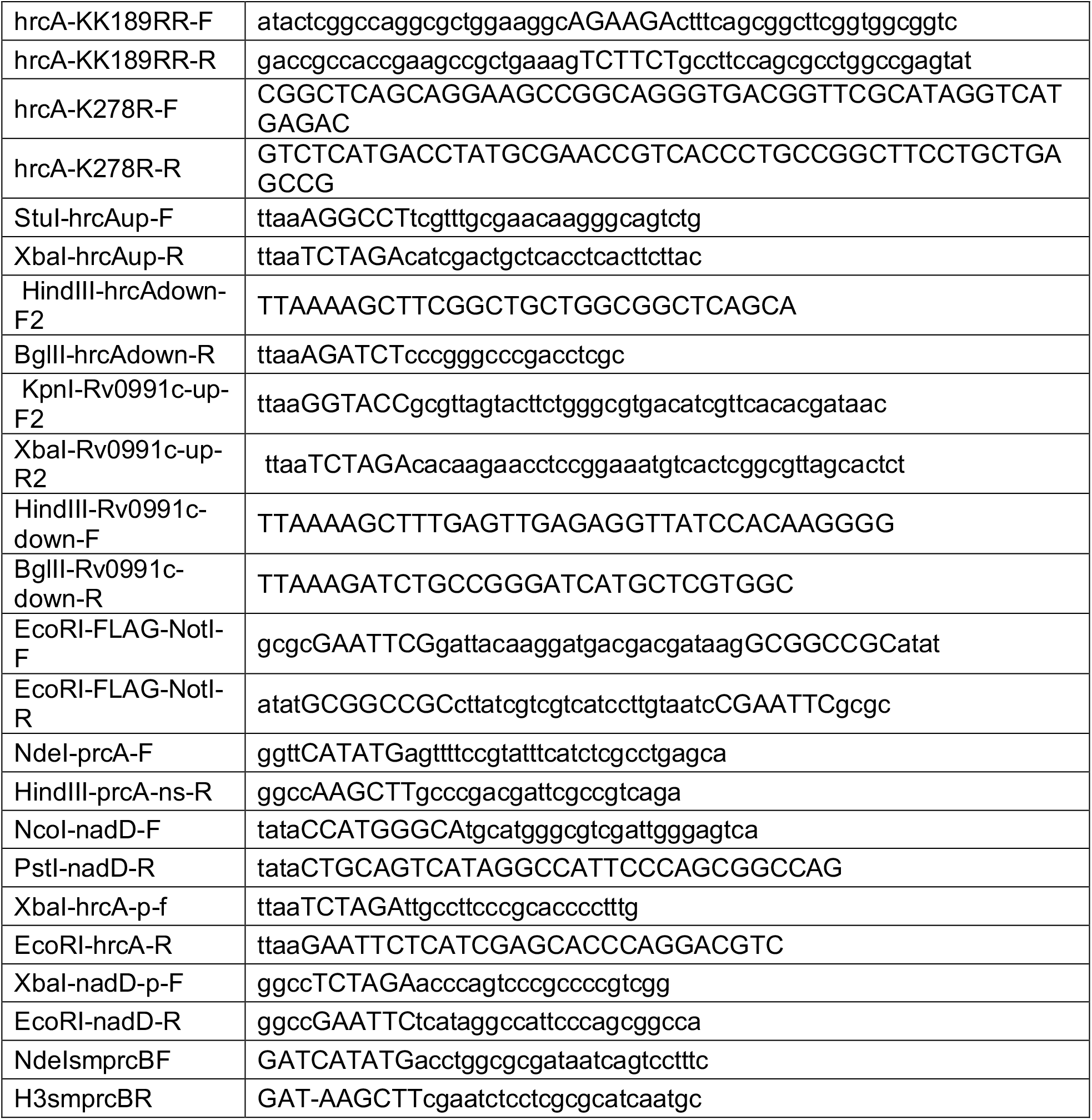
Strains, plasmids and primers used in this study.

### SUPPLEMENTARY FIGURE LEGENDS

**Figure S1.**
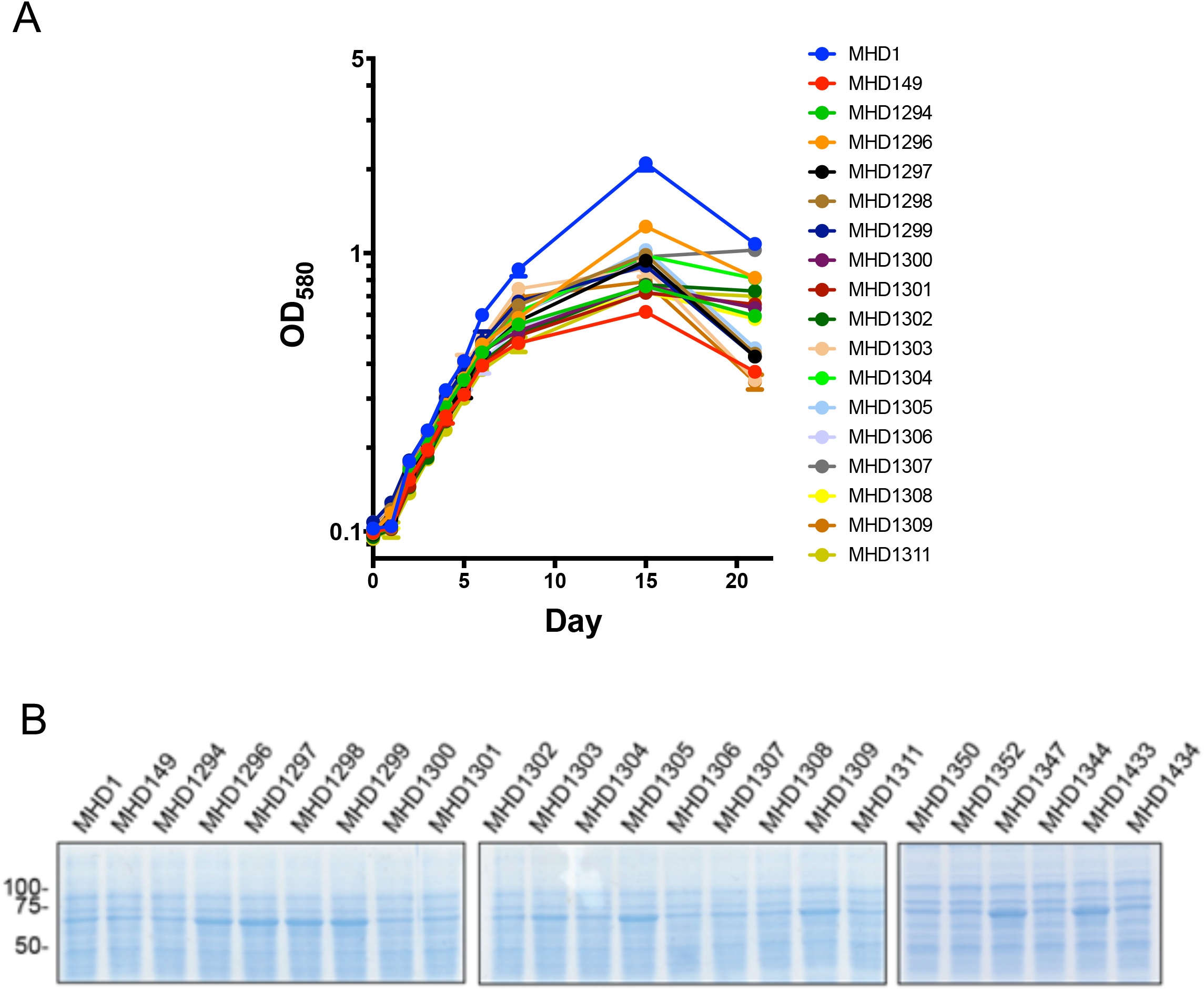
(A) Growth of indicated strains (see Table S1) in PB-nitrate. (B) Assessment of GroEL1/GroEL2 levels in suppressor mutants. Lysates were prepared from indicated strains (see Table S1) grown in 7H9; proteins were separated by SDS-PAGE and stained with Coomassie brilliant blue.

**Figure S2.**
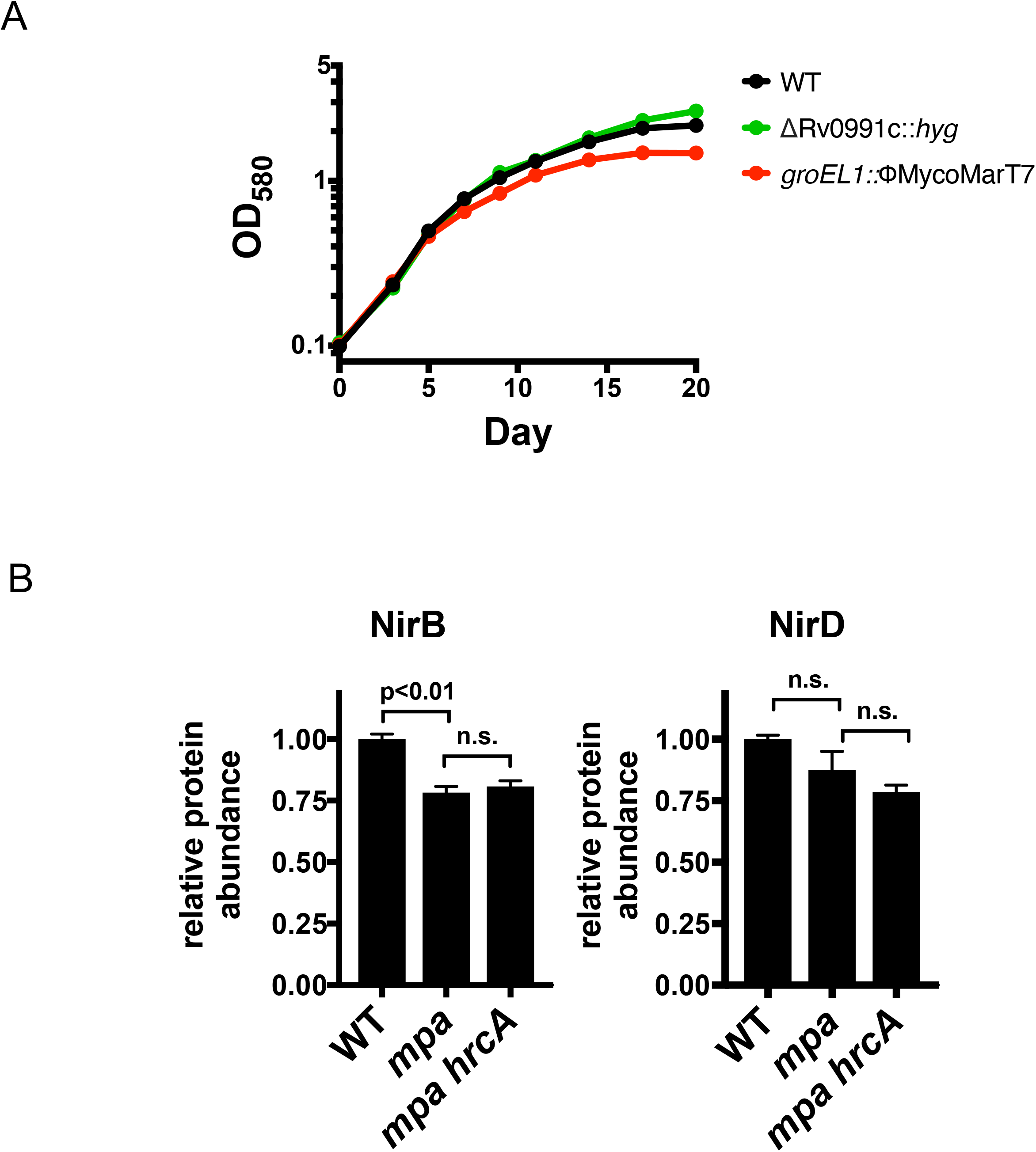
(A) Strains with mutations in Rv0991c (MHD1384) or *groEL1* (MHD15) grow in PB-nitrate. (B) NirB is less abundant in *mpa* (MHD149) and *mpa hrcA* (MHD1297) strains compared to WT, and NirD abundance follows a similar trend although the difference did not reach statistical significance. Data in (B) are adapted from Dataset S1.

**Figure S3.**
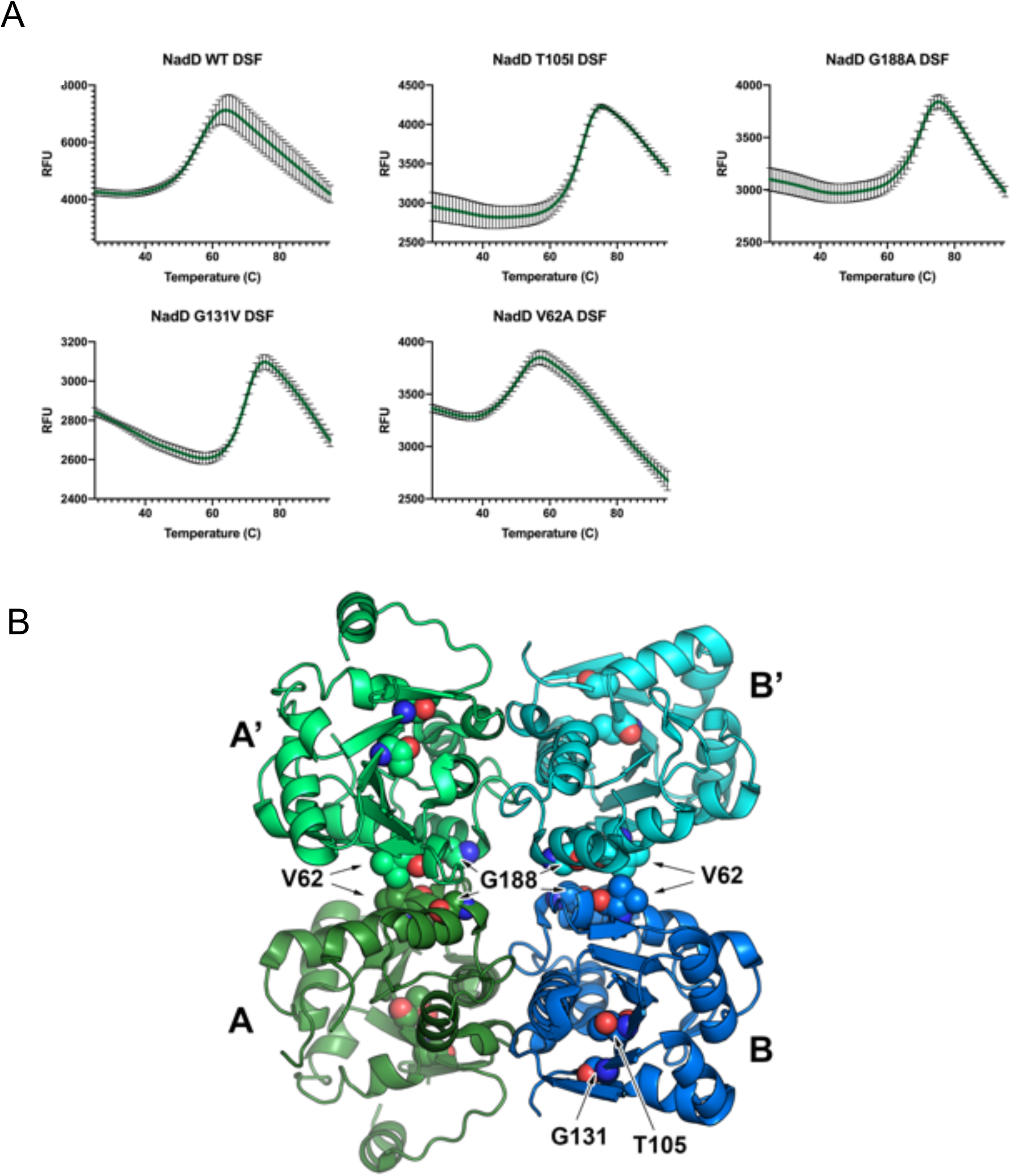
(A) Stability of purified WT NadD along with indicated NadD variants. Stability was determined by measuring protein melting temperature (T_m_) using a thermal shift assay (see Experimental Procedures in the main text). T_m_ values are summarized in Table 1. (B) Location of NadD mutations in the structure of *M. tuberculosis* NadD. A tetrameric structure of NadD (PDB: 4×0E) (7) is shown in cartoon representation. Mutated residues are shown as spheres. Chains A and B (which are present in the asymmetric unit) form a tetramer with the crystallographic symmetry chains A’ and B’.

### SUPPLEMENTARY EXPERIMENTAL PROCEDURES

#### Primers and plasmid construction

Table S1 contains a list of all primers and plasmids used in this study. Primers used for PCR amplification were purchased from Life Technologies. DNA was PCR-amplified using polymerases Phusion (New England Biolabs; NEB), Pfu (Agilent), or Taq (Qiagen) according to the manufacturers’ instructions. Plasmids encoding single- or double-lysine-to-arginine HrcA_TAP_ variants were constructed using overlap extension PCR (11). Amplified DNA was purified using the QIAquick Gel Extraction Kit (Qiagen). Restriction enzymes and T4 DNA ligase used for cloning were purchased from NEB, and cloning was performed in *E. coli* strain DH5a. Plasmids were purified from *E. coli* using the QIAprep Spin Miniprep Kit, and DNA was sequenced by GENEWIZ, Inc. to ensure the veracity of the cloned sequence. To generate pET24b(+)-*hrcA*-TAP by inserting the FLAG sequence directly upstream of His_6_, primers EcoRI-FLAG-NotI-F and EcoRI-FLAG-NotI-R were annealed and cloned into pET24b(+)-*hrcA*-his_6_. To construct pET24b(+)-*hrcA*_noK_-TAP, a plasmid harboring an *hrcA* allele with mutations in all six native lysine codons (*hrcA*_noK_) was constructed using gene synthesis (GENEWIZ, Inc.); *hrcA*_noK_ was subcloned into pET24b(+)-*hrcA-*TAP. To generate mutant variants of NadD, *nadD* mutant alleles were PCR-amplified from respective *M. tuberculosis* strains and cloned into the previously described construct for *M. tuberculosis* NadD expression in *E. coli* (7) (Table S1).

Deletion-disruption mutations in *hrcA* and Rv0991c (strains MHD1383 and MHD1384, respectively) were generated using allelic exchange plasmids to replace the respective genes with a hygromycin resistance cassette, as described in detail previously (8). For allelic exchange using pYUB854-*hrcA*-KO2, the last 77 codons of *hrcA* were maintained in order to preserve the promoter of the downstream gene, *dnaJ2* (Rv2373c). Successful allelic exchange in mutant strains was confirmed by PCR-amplifying and sequencing the deletion-disruption site from the chromosome.

Plasmids were transformed into *M. tuberculosis* by electroporation as previously described (12). Single-colony transformants were isolated on 7H11 with antibiotic selection.

#### Purification of recombinant NadD

To obtain NadD WT and mutant variants, Rosetta (DE3) *E. coli* competent cells were transformed with the relevant plasmids and grown to an OD_600_ of 1.5-1.7 at 37 °C with shaking. 400 µM Isopropyl β-D-thiogalactopyranoside (IPTG) was added and the cells were allowed to express for 4.5 hours at 37 °C. Because all four mutants displayed some level of cytotoxicity, the cells were grown to a higher density and allowed to express for a shorter time than is typical to minimize cell death while still providing acceptable protein yields. Pellets were obtained by centrifugation and cells were lysed with ThermoFisher’s bacterial protein extraction reagent (B-PER), lysozyme, and DNase at 25 °C for 1 hour. Protein was purified from filtered lysate by passage over a Ni-affinity column containing HisPur Superflow agarose (Thermo Fisher Scientific). Under these conditions, NadD co-purifies in a complex with NAD phosphate (NADP^+^). After washing with 20 mM Tris-HCl at pH 7.5, 300 mM sodium chloride, 10 mM imidazole, NADP^+^ was dissociated from NadD by three successive washes with buffer containing 20 mM Mg-ATP. Each wash was allowed a 20 min incubation. Mg-ATP was then removed by passage of three buffer washes with 15 min incubations before final elution using 20 mM Tris-HCl at pH 7.5, 300 mM sodium chloride, 250 mM imidazole. After overnight dialysis at 4°C to remove the imidazole, the protein was then further purified on a Superdex 200 10/300 Increase (GE Healthcare). The collected protein was then concentrated to 1 mg/ml using an Amicon Ultra-50 10 kD cutoff centrifugal filter unit before being flash frozen in liquid nitrogen.

#### Detailed immunoblotting procedures

Antibodies used in this study: FLAG M2 monoclonal antibody (Sigma-Aldrich) was used according to the manufacturer’s instructions; for Pup-Zur-His_6_ we used PentaHis Antibody (Qiagen); for GroEL2 immunoblots we used the NR-13655 monoclonal antibody to *M. tuberculosis* GroEL2 (BEI Resources) at a concentration of 1:1000 in 3% BSA; for Pup, we used an *M. tuberculosis* Pup-specific monoclonal antibody (13) at 1:1000 in 3% BSA; for DlaT immunoblots, DlaT antiserum (14) was used at 1:5,000 in 3% BSA; polyclonal rabbit antisera to InoI (15), Mpa (16), PrcB and PrcA were used at a 1:1000 dilution in 3% BSA. Secondary antibodies HRP-conjugated goat anti-rabbit IgG F(ab’)2 and HRP-conjugated anti-mouse IgG(H+L) were purchased from Thermo Fisher Scientific. All primary and secondary antibodies were made or diluted in 25 mM Tris-Cl/125 mM NaCl/0.05% Tween 20 buffer (TBST). Immunoblots were developed using SuperSignal West Pico PLUS chemiluminescent substrate (Thermo Fisher Scientific) and imaged using a Bio-Rad ChemiDoc system.

#### Detailed mass spectrometry procedures

Following preparation of lysates from *M. tuberculosis* as described in the main text, 150 µg of each protein lysate were reduced using dithiothreitol (5 μl of 0.2 M) for 1 h at 55 °C. The reduced cysteines were subsequently alkylated with iodoacetamide (5 μl of 0.5 M) for 45 min in the dark at room temperature. Next, 20 mM HEPES (pH 8.0) was added to dilute the urea concentration to 2 M. Protein lysates were digested with Trypsin (Promega) at a 100:1 (protein:enzyme) ratio overnight at room temperature. The pH of the digested protein lysates was lowered to pH < 3 using trifluoroacetic acid (TFA). The digested lysates were desalted using C18 solid-phase extraction (Sep-Pak, Waters). 40% acetonitrile (ACN) in 0.5% acetic acid followed by 80% ACN in 0.5% acetic acid was used to elute the desalted peptides. The peptide eluate was concentrated in a SpeedVac and stored at −80°C.

For tandem-mass-tag (TMT) labeling, the dried peptide mixture was re-suspended in 100 mM TEAB (pH 8.5) using a volume of 100 μl, and each sample was labeled with TMT reagent according to the manufacturer’s protocol (Thermo Fisher Scientific). In brief, each TMT reagent vial (0.8 mg) was dissolved in 41 μl of anhydrous ethanol and was added to each sample. The reaction was allowed to proceed for 60 min at room temperature and then quenched using 8 μl of 5% weight/volume hydroxylamine. The samples were combined at a 1:1 ratio and the pooled sample was subsequently desalted using SCX and SAX solid-phase extraction columns (Strata, Phenomenex) as described (17).

A 500 μg aliquot of pooled sample was fractionated using basic pH reverse-phase HPLC as previously described (18). Briefly, the sample was loaded onto a 4.6 mm × 250 mm Xbridge C18 column (Waters, 3.5 μm bead size) using an Agilent 1260 Infinity Bio-inert HPLC and separated over a 70 min linear gradient from 10 to 50% Buffer B in Buffer A at a flow rate of 0.5 ml/min (Buffer A = 10 mM ammonium formate, pH 10.0; Buffer B = 90% ACN, 10 mM ammonium formate, pH 10.0). A total of 40 fractions were collected throughout the gradient. The early, middle and late eluting fractions were concatenated and combined into 10 final fractions. The combined fractions were concentrated in the SpeedVac and stored at −80°C until further analysis.

For analysis by liquid chromatography with tandem mass spectrometry (LCMS/MS), an aliquot of each sample was loaded onto a trap column (Acclaim^®^ PepMap 100 pre-column, 75 μm × 2 cm, C18, 3 μm, 100 Å, Thermo Scientific) connected to an analytical column (EASY-Spray column, 50 m × 75 μm ID, PepMap RSLC C18, 2 μm, 100 Å, Thermo Scientific) using the autosampler of an Easy nLC 1000 (Thermo Scientific) with solvent A consisting of 2% ACN in 0.5% acetic acid and solvent B consisting of 80% ACN in 0.5% acetic acid. The peptide mixture was gradient eluted into the QExactive mass spectrometer (Thermo Scientific) using the following gradient: a 5%-23% solvent B in 100 min, 23%-34% solvent B in 20 min, 34%-56% solvent B in 10 min, followed by 56%-100% solvent B in 20 min. The full scan was acquired with a resolution of 70,000 (@ *m*/*z* 200), a target value of 1e6 and a maximum ion time of 120 ms. After each full scan 10 HCD MS/MS scans were acquired using the following parameters: resolution 35,000 (@ *m*/*z* 200), isolation window of 1.5 *m*/*z*, target value of 1e5, maximum ion time of 250 ms, normalized collision energy (NCE) of 30, and dynamic exclusion of 30 s.

Raw mass spectrometry data were processed using Proteome Discoverer 2.1. Proteins and peptides were searched against the Mycobacterium tuberculosis H37Rv proteome using the Byonic with a protein score cut-off of 300, using the following settings: oxidized methionine (M), and deamidation (NQ) were selected as variable modifications, and carbamidomethyl (C) as fixed modifications; precursor mass tolerance 10 ppm; fragment mass tolerance 0.02 Da. Proteins identified with less than two unique peptides were excluded from analysis. Bioinformatics analysis was performed with Perseus and Microsoft Excel. Student’s t-test using Benjamini-Hochberg FDR cutoff was then used to identify proteins with differential abundance between bacterial strains.

